# Improved nucleoside (2’-deoxy)ribosyltransferases maximize enzyme promiscuity while maintaining catalytic efficiency

**DOI:** 10.1101/2025.03.05.641663

**Authors:** Peijun (Gary) Tang, Greice M. Zickuhr, Alison L. Dickson, Christopher J. Harding, Suneeta Devi, Tomas Lebl, David J. Harrison, Rafael G. da Silva, Clarissa M. Czekster

## Abstract

Nucleoside analogues have been extensively used to treat viral and bacterial infections and cancer for the past 60 years. However, their chemical synthesis is complex and often requires multiple steps and a dedicated synthetic route for every new nucleoside to be produced. Wild type nucleoside 2′-deoxyribosyltransferase enzymes are promising for biocatalysis. Guided by the structure of the enzyme from the thermophilic organism *Chroococcidiopsis thermalis* PCC 7203 (*Ct*NDT) bound to the ribonucleoside analogue Immucillin-H, we designed mutants of *Ct*NDT and the psychrotolerant *Bacillus psychrosaccharolyticus* (*Bp*NDT) to improve catalytic efficiency with 3′-deoxynucleosides and ribonucleosides, while maintaining nucleobase promiscuity to generate over 100 distinct nucleoside products. Enhanced catalytic efficiency towards ribonucleosides and 3′-deoxyribonucleosides occurred via gains in turnover rate, rather than improved substrate binding. We determined crystal structures of two engineered variants as well as kinetic parameters with different substrates, unveiling molecular details underlying their expanded substrate scope. Our rational approach generated robust enzymes and a roadmap for reaction conditions applicable to a wide variety of substrates.

**Insert Table of Contents artwork here:** 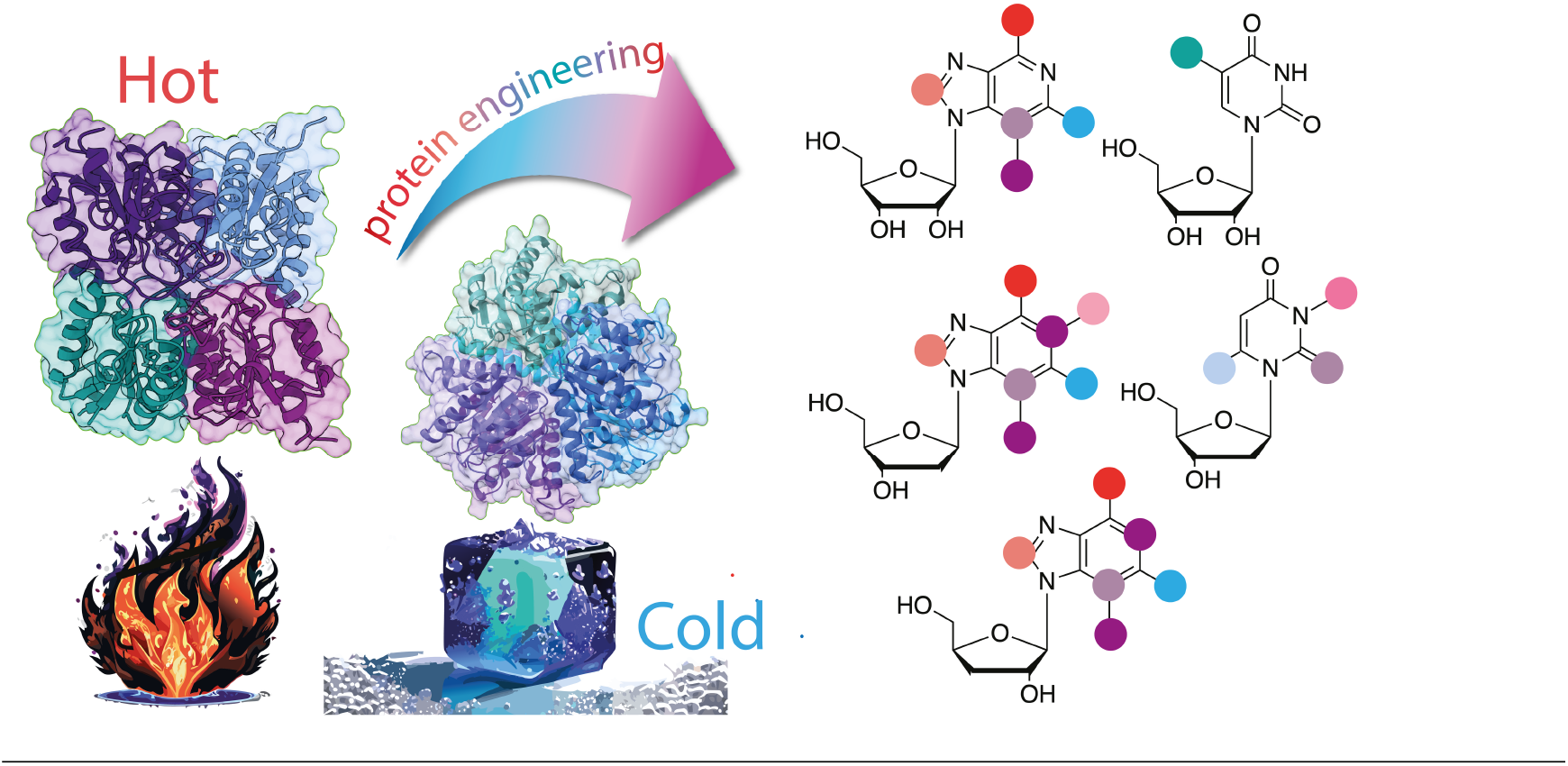

---

Nucleoside analogues (NAs) are used to treat cancer, viral and bacterial infections.^1^ They are challenging to produce synthetically, and enzymatic routes are an alternative to access novel analogues.^2^ Merck’s biocatalytic synthesis of the islatravir demonstrated the feasibility of an *in vitro* fully biocatalytic cascade for the synthesis of an anti-HIV NA.^3^ Moreover, incorporating nucleoside 2′-deoxyribosyltransferase (NDT) enzymes for the fully biocatalytic production of the NA anti-SARS-CoV-2 drug Molnupiravir has been proposed.^4^ Novel nucleoside and nucleotide analogues chemically synthesized for targeting human cancers and bypassing resistance to common treatments have been proposed, demonstrating scope for future work on nucleoside development.^5^ Importantly, recent work demonstrated the versatility of NDTs and other enzymes from nucleoside salvage pathways to generate new NAs.^6-7^

Enzymes that display broad substrate scope and the capacity to catalyse more than one type of reaction are desirable, albeit less explored.^8^ Here we employed two nucleoside 2′-deoxyribosyltransferases, the enzyme from the thermophilic organism *Chroococcidiopsis thermalis* PCC 7203 (*Ct*NDT) and the enzyme from the psychrotolerant organism *Bacillus psychrosaccharolyticus* (*Bp*NDT) to produce a series of over 40 NAs, several of which unprecedented (Figure 1). Furthermore, we explored the effects of reaction conditions, including pH, temperature, substrate concentrations and usage of a coupled enzyme in efforts to drive the reaction equilibrium towards desired products and decrease nucleoside hydrolysis uncoupled from nucleobase transfer. We established a workflow to optimize reaction conditions and determine factors affecting reaction yield, with implications for others working in nucleoside biocatalysis. We also determined the structure of a double mutant with improved catalytic efficiency towards 3′-deoxynucleosides and ribonucleosides, establishing a path to guide future NDT engineering which relies on modulating steric effects and electrostatics surrounding a conserved and essential catalytic glutamate residue.

**Figure 1:**
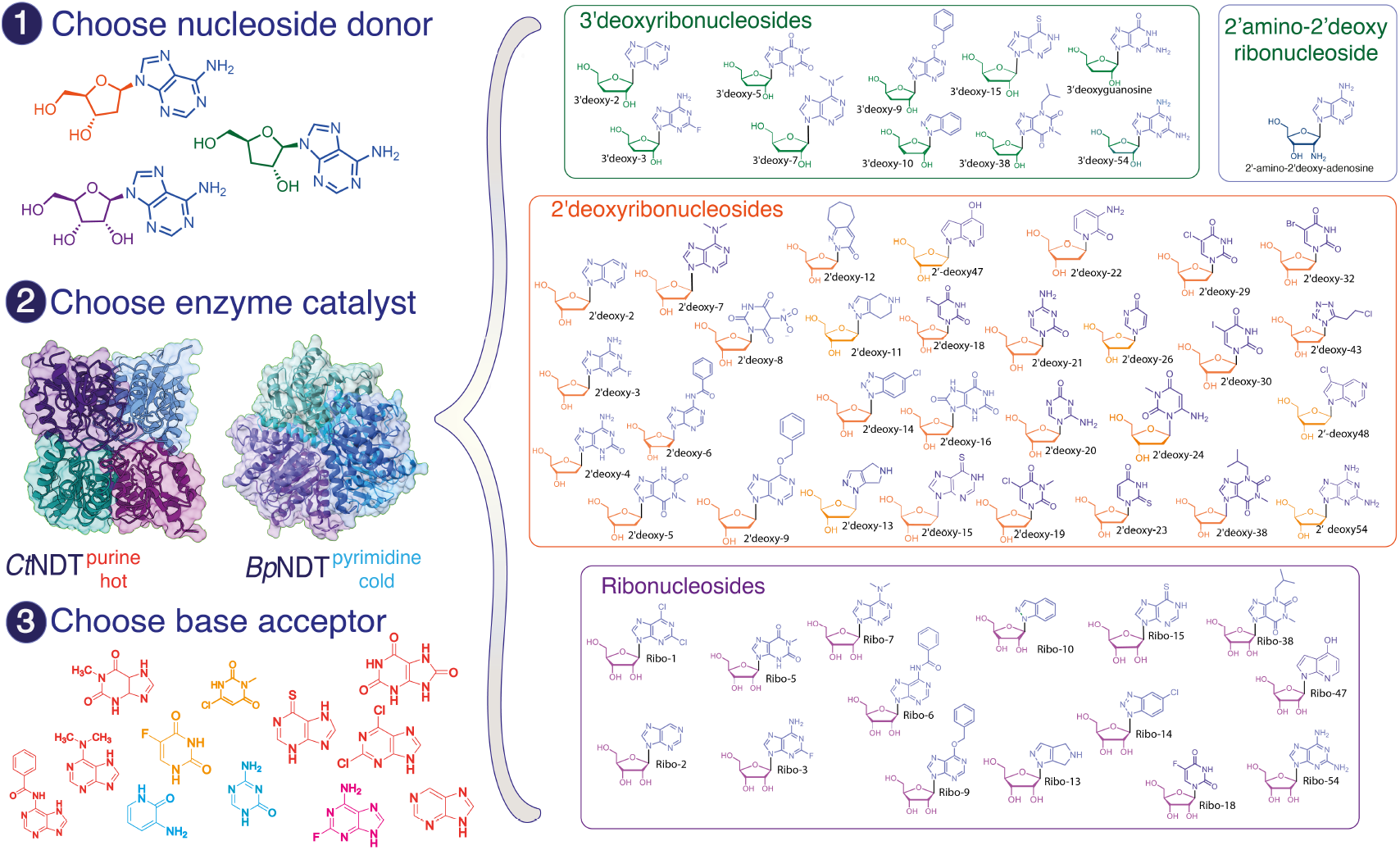
Nucleoside and nucleobase substrate scope and generation of novel nucleosides using *Ct*NDT or *Bp*NDT. Left: 1) A nucleoside donor is chosen between a ribonucleoside, a 2′−deoxy-ribo-nucleoside or 3′−deoxyribonucleoside. 2) An enzyme catalyst is chosen based on nucleoside donor identity and reaction conditions since *Ct*NDT operates at high temperatures, while *Bp*NDT operates at low temperatures. 3) A nucleobase acceptor is also chosen. Right: Nucleosides produced with a combination of *Ct*NDT or *Bp*NDT variants. HRMS data for each product generated are on Figure S1 and Table S2.

Inspired by the structures of *Trypanossoma brucei* (*Tb*NDT)^9^, *Bp*NDT^10^ and our recent work on *Ct*NDT^11^, we performed structure-based enzyme engineering to produce modified *Ct*NDT enzyme variants with expanded substrate scope. According to reports with other NDT enzymes^7, 12^, we first explored the natural promiscuity of *Ct*NDT and *Bp*NDT. *Ct*NDT has a preference for purine-based 2′-deoxynucleosides, while *Bp*NDT prefers pyrimidine-based 2′-deoxynucleosides (Figure S1). Both enzymes can utilize a vast array of non-canonical nucleobases as substrates, with *Ct*NDT possessing a broader substrate scope. Figure 1 depicts the NAs produced here, employing wild type and enzyme variants discussed below. Figure S1 and Table S2 show more details on substrates tested and products obtained. Importantly, wild type *Ct*NDT can utilize ribonucleoside and 3′-deoxynucleoside substrates, albeit with lower efficiency, demonstrating less strict selection of 2′-deoxynucleosides, and a possible strategy to produce 3′-deoxynucleosides.^7, 13^ As proof of concept, we determined cordycepin (3′-deoxyadenosine), a natural product currently employed to treat various types of cancers, can be used as a substrate by *Ct*NDT.^14^ This enabled the generation of other 3′-deoxynucleoside derivatives using *Ct*NDT variants. Importantly, bulky nucleobases such as 6-(benzyloxy)-9H-purine were accepted as substrates, as well as 3-aminopyridin-2(1H)-one, which is not typically considered a nucleobase. *Ct*NDT has an average sized substrate binding pocket in comparison to other dNDT enzymes (Figure S3), with a solvent accessible volume of ∼ 150 Å^3^, smaller than *Lactobacillus leichmannii* (LlNDT, ∼ 170 Å^3^), which was also shown to accept many different nucleobases and 2′-deoxynucleoside as substrates.^7, 15^ The synthesis of purine nucleosides with bulkier, expanded bases using traditional chemical synthesis has been challenging, despite some showing promise as anticancer and antiviral compounds^16^.

An asparagine close to the 3′-OH group of ribonucleosides has been proposed as a “gatekeeping residue” controlling acceptance of ribonucleoside substrates, and it is replaced by an aspartate on other enzymes that do not utilize ribonucleosides as substrates. *Ct*NDT lacks this “gatekeeping” asparagine (N53 on *Tb*NDT is replaced by D62 on *Ct*NDT). Recent work has exploited a double mutant (residues equivalent to Y7F/D62N in *Ct*NDT) in the enzyme from *L. leichmannii*^*15*^ to improve acceptance of ribonucleoside substrates, but an improvement in product conversion was not observed for this mutant. A comparison between the structures of *Ct*NDT mutants and *Lh*NDT and *Ll*NDT is shown on Figure S3. The mutant *Ct*NDT_D62N_ does not display improved kinetic parameters with ribonucleoside substrates (Figure 3), hence a different nucleoside selection strategy is likely taking place.

Prior work carried out the mutation of a tyrosine to phenylalanine in the vicinity of the sugar binding pocket of NDT enzymes to improve ribonucleoside substrate utilization.^17^ We have previously shown that this tyrosine residue in *Ct*NDT (Y7) does not directly interact with the 2′-OH group, and instead positions the catalytic E88 for reaction.^11^ This tyrosine residue is conserved in NDT enzymes, and further explored below.

Crucial to our engineering efforts, as well as future applications of NDTs as biocatalysts, we obtained crystal structures of engineered mutants bound to ribonucleoside analogues, providing atomic-level detail into how changes in the ribosyl-binding pocket accommodate these modified substrates (Figure 2 and Tables S4 and S5). To understand the interaction between *Ct*NDT and ribonucleosides, we co-crystallized the variant *Ct*NDT_Y7F_, which possesses higher catalytic efficiency with ribonucleoside substrates than *Ct*NDT_WT_ with the nucleoside analogue Immucillin-H (ImmH). This analogue is a purine nucleoside phosphorylase inhibitor, rationally designed to mimic the transition state of the reaction catalysed by that enzyme. Importantly, for *Ct*NDT it is a non-hydrolysable substrate analogue containing hydroxyl groups on positions 2′ and 3′. Figure 2a compares the structures of *Ct*NDT_WT_ and *Ct*NDT_Y7F_, depicting key residues participating in the reaction and substrate selection. A flexible loop that acts as a “lid” and could potentially allow bulkier substrates to be used was observed in an “open” or “closed” conformation (Figure 2a and Figure S2). Figure S3 depicts differences in the normalized B-factors for this flexible loop and ligands, as only 2 molecules of ImmH were placed in the *Ct*NDT_Y7F_ tetramer, with less defined electron density in the nucleobase moiety of ImmH, while two other monomers remained unoccupied.

**Figure 2:**
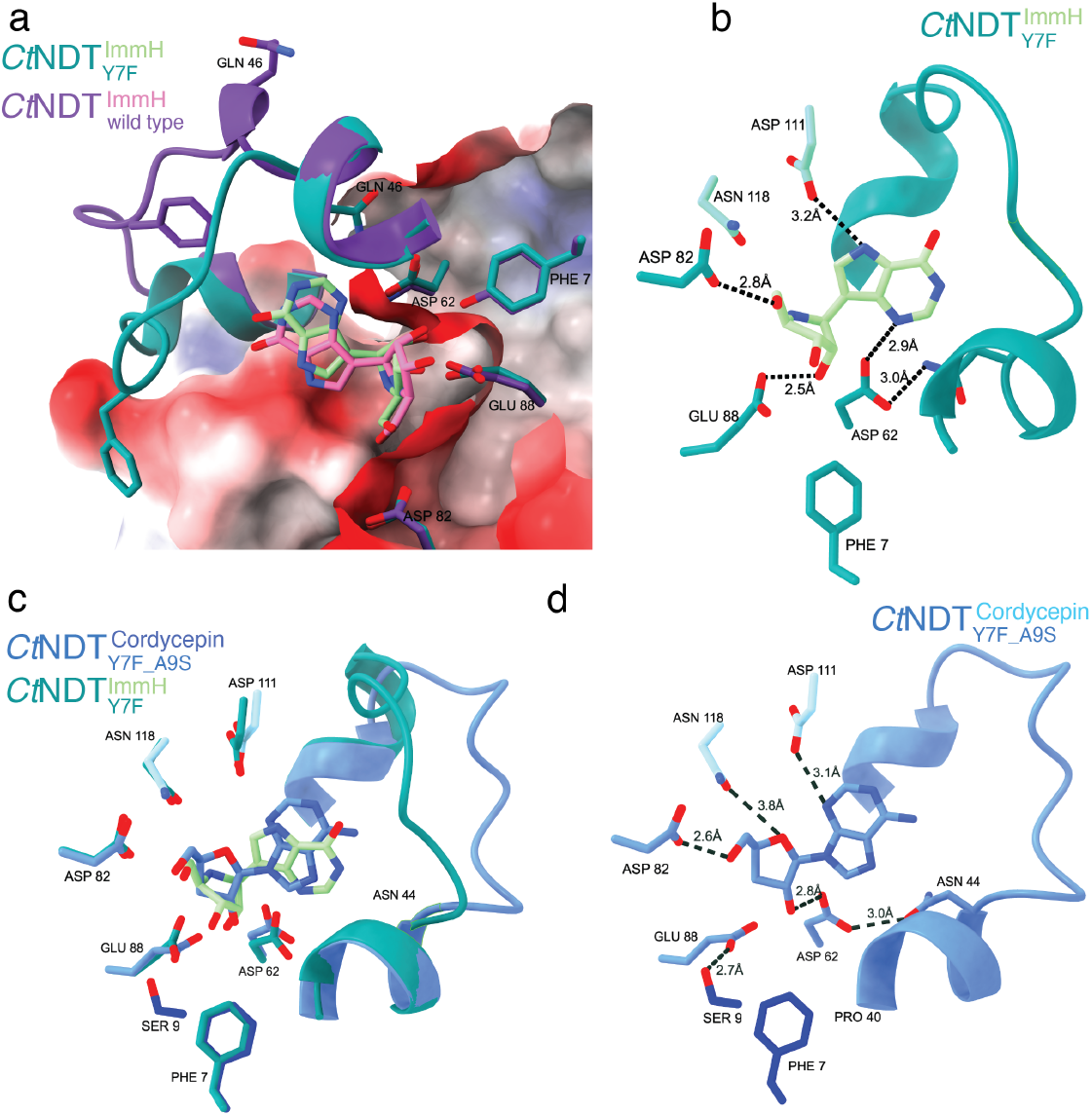
Structural features of enzyme variants characterized here. a) Overlay of structures of wild type *Ct*NDT (pdb 8PQP, purple/pink) and *Ct*NDT_Y7F_ bound to ImmucillinH (ImmH, teal/green). Surface colored to depict electrostatic potential. Loop covering active site and Gln46 occupy different positions in the open and closed conformation shown. b) Interactions between *Ct*NDT_Y7F_ and ImmH. c) Overlay of structures of *Ct*NDT_Y7F_ (teal/green) and *Ct*NDT_Y7F_A9S_ (blue) bound to ImmH and cordycepin, respectively). d) Details of interactions between *Ct*NDT_Y7F_A9S_ and cordycepin. Key distances are shown, and residues mutated (Tyr7, Ser9) are shown in dark blue. Loop covering the active site depicted for reference. In b-d residues 111 and 118 are from a neighbouring subunit.

**Figure 3:**
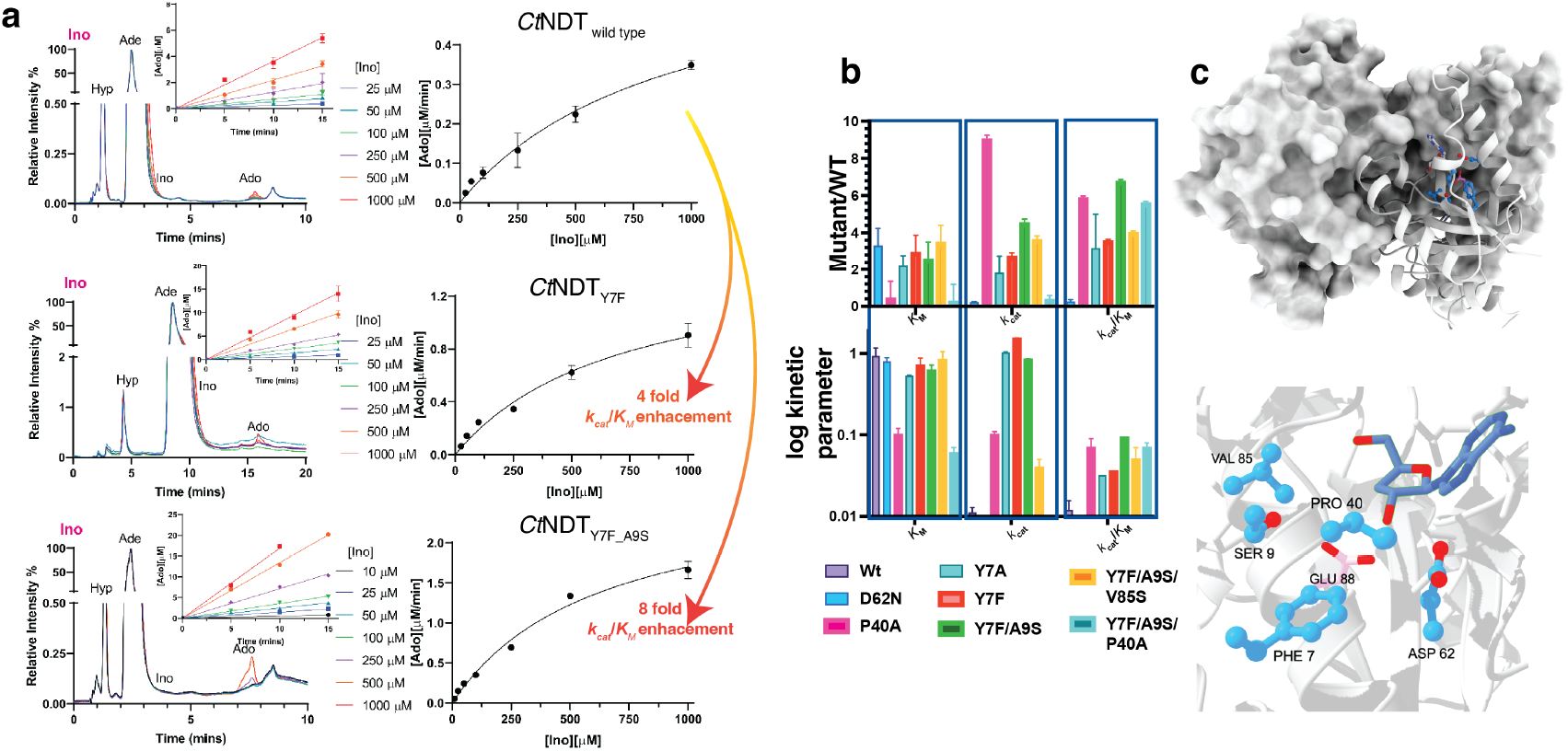
Kinetic characterization of enzyme variants with Ino and Ade as substrates: a) Left: raw HPLC data used to obtain initial rates at different substrate concentrations (inset) and generate Michaelis-Menten plots (right). b) Summary of kinetic parameters obtained for each enzyme variant Experiments were conducted in triplicate and data are shown as average and spread error derived from standard error of fits. c) Residues mutated in variants under study. On the top the overall tetramer is shown, and in the bottom a zoomed-in image of the substrate (cordycepin, dark blue) and mutated residues (marine blue), depicted here from *Ct*NDT_Y7F-A9S_.

No major differences are present in the protein backbone, but the wild-type enzyme binds ImmH in a distorted conformation in relation to the ribose ring, bringing the 2′-OH farther away from D62 and closer to Y7, likely to avoid clashes (Figure 2a), which does not occur in the *Ct*NDT_Y7F_ mutant. A DALI search indicates the closest NDT homologues in the PDB are the proteins from *Trypanosoma cruzi* (pdb 2f67^9^, rmsd 0.85 Å), *Leishmania mexicana* (6qai^18^, rmsd 0.87 Å), and *Lactobacillus helveticus* (1s2g^19^, rmsd 0.89 Å – henceforth referred to as *Lh*NDT). *Lh*NDT complex with 2′-deoxyadenosine allows a comparison between residues interacting with base and nucleoside moieties (Figure S3a). Following our interaction map with ImmH, a ribonucleoside analogue, we designed mutants targeting residues surrounding the 2′-OH group, aimed at further tailoring substrate selection towards modified sugars. Our rationale was centred in (*i*) opening up the substrate binding pocket by mutating P40 and Y7 and (*ii*) and exploring additional hydrogen bonding interactions to better position a 2′-OH group by mutating A9 and V85. Single mutants *Ct*NDT_A9S_, *Ct*NDT_V85S_, *Ct*NDT_Y7A_, *Ct*NDT_P40A_, double mutant *Ct*NDT_Y7F-A9S_ and triple mutants *Ct*NDT_Y7F-A9S-V85_ and *Ct*NDT_Y7F-A9S-P40A_ were evaluated with inosine as sugar donor and adenine as sugar acceptor. These mutants had modest changes on *K*_M-inosine_, and the most pronounced effects were driven by *k*_cat_ (Figure 3 and Figure S4). The *K*_D_ for Immucillin-H for the double mutant *Ct*NDT_Y7F-A9S_ is 590 μM, 7.5 times higher than the one determined for the wild-type protein (Figure S2), and close to the *K*_M_ for inosine (620 μM, Table S3), therefore binding in a substrate-like manner. Figure 2c shows the complex structure of *Ct*NDT_Y7F-A9S_ and Cordycepin. A potential additional interaction between S9 and the catalytic E88 (distance 2.5 Å) is formed. This could be important to compensate for the loss in positioning previously conferred by Y7, mutated to a phenylalanine in this variant. In agreement with this, *Ct*NDT_Y7F-A9S_ showed an improvement on *k*_cat_/*K*_M-inosine_ of eight-fold in comparison to wild type, and a two-fold improvement towards *Ct*NDT_Y7F_.

Since pyrimidines were not efficient substrates for *Ct*NDT, we turned to *Bp*NDT as it was previously shown to use 2’-deoxynucleoside pyrimidine substrates.^10, 20^ In our enzymatic assays, analogues of 2’-deoxyuridine were produced. Given 5-fluoro-2′-deoxyuridine (Floxuridin) and capecitabine are FDA approved to treat different cancers, new analogues are desirable. We generated the mutant *Bp*NDT_Y5F_, which is equivalent to *Ct*NDT_Y7F_ since *Bp*NDT_WT_ cannot utilize ribonucleosides as substrates. *Bp*NDT_Y5F_ uses guanosine and uracil as substrates to produce 5-fluorouridine as a product (Figure S1, Table S2). Cordycepin and clofarabine were not substrates for *Bp*NDT_WT_ or *Bp*NDT_Y5F_, demonstrating this enzyme has a narrower substrate scope than *Ct*NDT. Both *Bp*NDT^21^ and *Ct*NDT^22^ have been previously immobilized, further illustrating the potential of these enzymes in nucleoside production.

To explore the applicability of employing *Ct*NDT, *Ct*NDT_Y7F-A9S_ and *Bp*NDT to produce novel compounds, we developed a substrate scope and reaction condition testing matrix as summarized on Figure 1 and detailed on Figure 4. Depending on the enzyme variant, an optimal nucleoside is chosen to act as “sugar donor”, while nucleobase is varied. When sufficient quantities of nucleobase are available, increasing the ratio nucleobase/nucleoside can increase reaction yields to up to 99% when considering the limiting substrate. Other factors influence reaction yields, including pH, temperature and reaction times (Figure S6), and we hypothesize this is due to reaction kinetics x reaction equilibrium when different substrates are employed. Similar to what is observed with nucleoside phosphorylases^23^, we hypothesized that in cases where 2′-deoxyinosine acts as nucleoside sugar donor, adding xanthine oxidase could increase product formation specially in earlier time points. However, uric acid is a substrate for *Ct*NDT with a *k*_cat_/*K*_M-uric_acid =_ 0.22 mM^-1^s^-1^ in the same range as observed for guanine for example (0.33 mM^-1^s^-1^), and therefore over time it is consumed as a nucleobase substrate,shifting the equilibrium towards initial conditions. In the second half reaction for dNDTs, after the ribosyl intermediate is formed there is competition between water and the incoming nucleobase for product formation leading to either nucleoside hydrolysis or formation of a new nucleoside product. We determined the hydrolysis rate (Figure S5e), and decrease in yield over time could be due to nucleoside hydrolysis, although the identity and concentration of nucleobase substrate will also influence yield including when uric acid, xanthine and hypoxanthine which have similar *k*_cat_/*K*_M_ values and are equally competent substrates are present in the reaction mixture. Furthermore, commercially available xanthine oxidase is not active for longer than 2-4h in some reaction conditions (depending on pH and temperature), and therefore unlikely to be a useful strategy at longer time points. All these factors render the use of xanthine oxidase with *Ct*NDT impractical.

**Figure 4:**
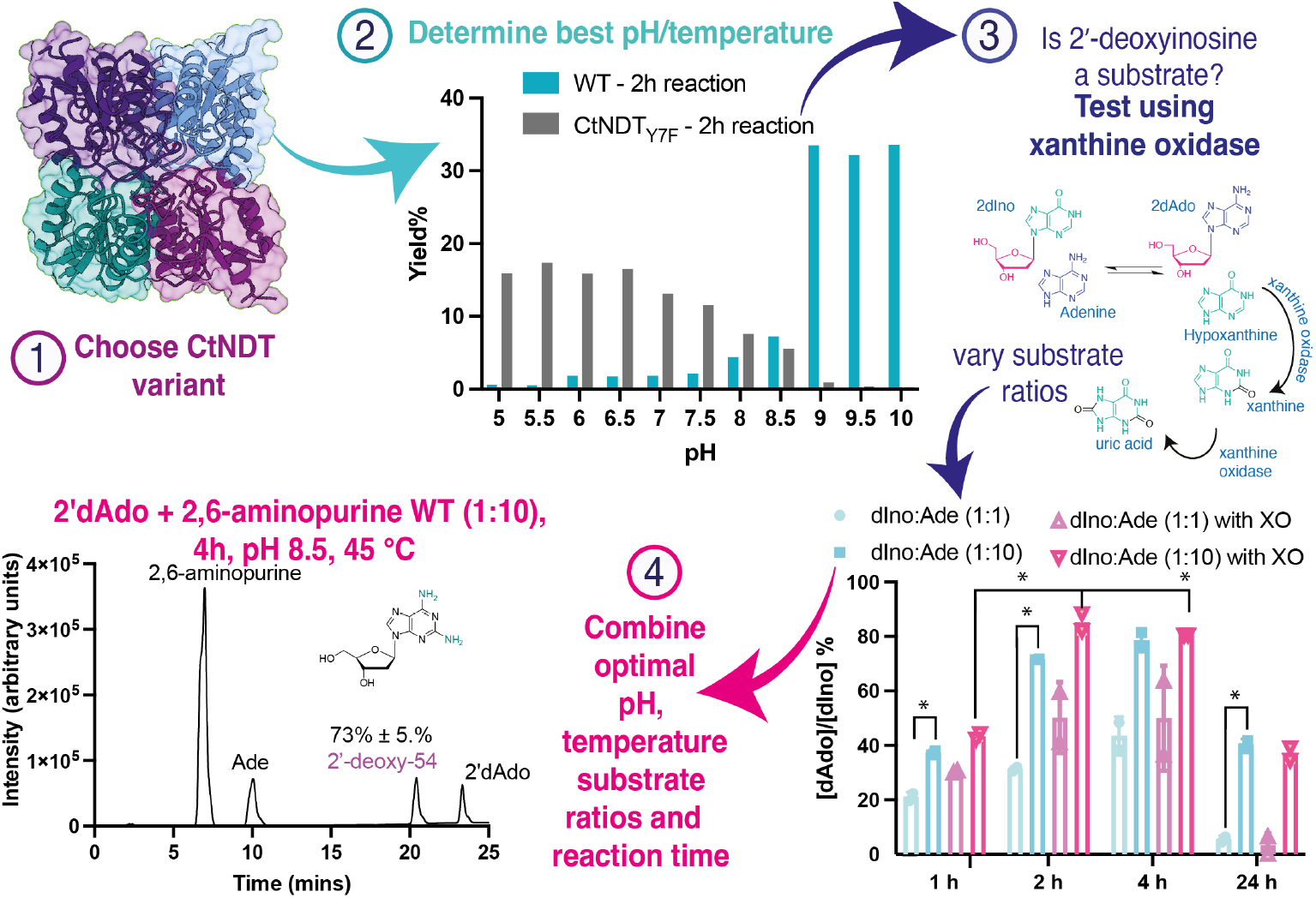
Steps to optimize reaction conditions and determine kinetic parameters, while also improving reaction yield. Step 1: choose a CtNDT variant to test (depending on the nucleoside substrate as wild type, *Ct*NDT_Y7F_ and *Ct*NDT_Y7F-A9S_ were shown to possess different preferences); step 2: determine the optimal pH and temperature to monitor the reaction taking into account enzyme stability and substrates solubility; step 3: test whether xanthine oxidase can improve reaction yield at different ratios of substrates (graph shows significant differences in yield with p<0.0332, in this case reaction times significantly affect yield, but not the presence of xanthine oxidase); step 4: combine optimal substrate ratios (nucleoside and nucleobase), reaction time, pH and temperature to scale up reaction. Figure S6 shows more details on reaction yield.

Applying our condition matrix, we determined yield and developed a purification protocol for 2-fluoro-3′-deoxyadenosine (or 2F-3′d-Ado or 2F-cordycepin, compound 3’-deoxy-3), and 2-fluoroadenosine (compound ribo-3) using both substrates at a 1:1 ratio, and producing N-(9H-purin-6-yl)benzamide (compound ribo-6) and 2′-deoxy-2-amino-adenosine (compound 2’-deoxy-54) using a 10:1 nucleobase to nucleoside substrate ratio. Yields were 13%, 80%, 78%, and 73% respectively. 2F-cordycepin was previously synthesized with 2% overall final yield starting from adenosine and shown to be a potent anti-trypanosomal compound^24^. NDTs can also act as nucleoside hydrolases in the absence or under limiting concentrations of the nucleobase substrate in the second half reaction, and we evaluated nucleoside substrate hydrolysis to have a variable effect on reaction yield (from no hydrolysis to up to 23% hydrolysis of 3’-deoxy-adenosine when *Ct*NDT_Y7F-A9S_ was employed (Figure S5).

In summary, we engineered NDT enzymes to broaden substrate scope and generate over 40 nucleoside analogues, several of which currently have no synthetic route proposed. We established a workflow to determine optimal reaction conditions and generated a mutant (*Ct*NDT_Y7F-A9S_) with altered nucleoside substrate specificity towards 3’-deoxynucleosides and ribonucleoside substrates.

## ASSOCIATED CONTENT

### Supporting Information

The Supporting Information is available free of charge on the ACS Publications website. Materials and methods, raw data for LC-MS and HPLC traces, intact mass for proteins employed here, saturation curves and additional figures and tables are available.

## AUTHOR INFORMATION

### Author Contributions

C.M. Czekster, R.G. da Silva, and D.J. Harrison conceptualized the project and discussed data, discussed and proofread the article;

T. Lebl provided support with compound characterisation; P. Tang, A. Dickson, C.J. Harding, G.M. Zickuhr, and S. Devi performed experiments, wrote and proofread the manuscript.

### Notes

D.J.H. is part-time employed by NuCana plc. A.D. is employed by Compass Pathways.

## ACKNOWLEDGMENT

We thank the mass spectrometry facility in St Andrews for support in mass spectrometry.

## ABBREVIATIONS

PCC 7203: 2′-deoxyribosyltransferase from *Chroococcidiopsis thermalis*
(*Ct*NDT, Uniprot K9TVX3): *Bacillus psychrosaccharolyticus* (*Bp*NDT, Uniprot A0A3G5BRZ6
(ImmH): Immucillin-H.

## PDB Accession codes

9EMX - *Ct*NDT_Y7F-A9S_ bound to Cordycepin, and 9EMW -

*Ct*NDT_Y7F_ bound to ImmH.

