## Supporting information for "Improved nucleoside (2’-deoxy)ribosyltransferases maximize enzyme promiscuity while maintaining catalytic efficiency"

### Peijun (Gary) Tang[a], Greice M. Zickuhr[b], Alison L. Dickson[b], Christopher J. Harding[c], Suneeta Devi[d], Tomas Lebl[e], David J. Harrison[b,f], Rafael G. da Silva[a] and Clarissa M. Czekster [a,*]

[a]School of Biology, Biomedical Sciences Research Complex, University of St Andrews, St Andrews, Fife KY16 9ST, United Kingdom; [b]School of Medicine, University of St Andrews, North Haugh, St Andrews, KY16 9TF, UK; [c] Institute of Infection, Veterinary and Ecological Sciences, University of Liverpool, L69 3BX, UK; [d] School of Physical Sciences, University of Liverpool, L69 7ZF, UK; [e]School of Chemistry and Biomedical Sciences Research Complex, University of St Andrews and EaStCHEM, North Haugh, St Andrews, Fife, KY16 9ST, United Kingdom; [f]NuCana Plc, Edinburgh, EH12 9DT, United Kingdom

#### Materials and methods

##### Abbreviations

2′-deoxyadenosine (2′-dAdo), 2′-deoxyguanosine (2′-dGuo), 2′-deoxyinosine (2′-dIno), 2´-dUridine (2′-dUrd), 2´-dCytidine (2′-dCyd), 2′-dThymidine (2′-dThd), Adenine (Ade), Guanine (Gua), Hypoxanthine (Hyp), Cytidine (Cyt), Adenosine (Ado), Guanosine (Guo), Inosine (Ino), Uracil (Ura), Uric acid (UricA), liquid-chromatography-mass spectrometry (LC-MS), polyethylene glycol (PEG), Immucillin H (ImmH).

##### Materials

The synthetic gene was ordered from Integrated DNA Technologies (IDT). The chemicals and reagents were provided by Fluorochem, Merck and Fisher Scientific. Isocytidine was provided by Nucana plc.

##### Additional Methods

##### Cloning, expression and purification of *Ct*NDT mutants

The original synthetic gene encoding the *Chroococcidiopsis thermalis* NDT (Uniprot code: K9TVX3) and *Bacillus psychrosaccharolyticus* NDT (Uniprot code: A0A3G5BRZ6) were cloned into a pJ411 (*Ct*NDT) or pJ414 (*Bp*NDT) expression plasmid with a cleavable 6-histidine tag at N-terminus. Single mutants *Ct*NDT$_{A9S}$, *Ct*NDT$_{V85S}$, *Ct*NDT$_{Y7A}$, *Ct*NDT$_{P40A}$, double mutant *Ct*NDT$_{Y7F\text{-}A9S}$ and triple mutants *Ct*NDT$_{Y7F\text{-}A9S\text{-}V85}$ and *Ct*NDT$_{Y7F\text{-}A9S\text{-}P40A}$, as well as *Bp*NDT$_{Y5F}$ were cloned with primers listed on Table S1 (generated using NEBaseChanger - New England Biolabs Inc., Massachusetts, USA). Mutagenesis was carried out using the KLD (Kinase–Ligase–DpnI) enzyme mix (M0554S, New England Biolabs Inc., Massachusetts, USA) to remove the DNA template and ligate the new amplicon, followed by transformation into NEB 5alpha cells. All mutations were confirmed by sequencing (Eurofins).

The expression and purification protocols for the mutants are the same as previously described[1], except *Bp*NDT remained in the histidine tag due to protein instability after tag removal[2]. In summary, the purification protocol was as follows:

The synthetic gene encoding the *Chroococcidiopsis thermalis* NDT (Uniprot code: K9TVX3) was ordered as a Gblock (Integrated DNA Technologies) and cloned into a pJ411 expression plasmid with a cleavable 6-histidine tag at the N-terminus by Gibson assembly. Construct design was carried out using NEBuilder (New England Biolabs), following the cloning protocol suggested to amplify the plasmid pJ411 backbone as well as the GBlock to generate an overlap of 20 bp between sequences. Following cloning, gene sequence was confirmed by sequencing (Eurofins). pJ411::*Ct*NDT and pJ414::*Bp*NDT were transformed into *E.coli* BL21 (DE3) cells for overexpression in LB medium with 50 mg/mL kanamycin or 100 mg/mL ampicillin at 37°C with shaking at 180 rpm shaking until cells reached $OD_{600}$ = 0.8. Protein expression was induced by addition of 0.5 mM of IPTG overnight at 16°C with shaking at 180 rpm. Cells were harvested by centrifugation at 12,000 g for 20 min, resuspended in wash buffer (50 mM MES, 250 mM NaCl, 30 mM imidazole, pH 6.5) and lysed using a cell homogenizer (Constant Systems). Following centrifugation for 30 min at 51,000 *g* at 4°C, the supernatant was filtered with a 0.8 mm filter to remove particulates and loaded onto a 5 mL HisTrap column pre-equilibrated with wash buffer. The column was washed with 10 column volumes of wash buffer, and *Ct*NDT and *Bp*NDT were both eluted with 50 mM MES, 250 mM NaCl, 500 mM Imidazole, pH 6.5. After the first HisTrap column, *Bp*NDT and its mutant *Bp*$NDT_{Y5F}$ were dialysed overnight to remove imidazole, then concentrated and stored at -80 °C.

Fractions containing *Ct*NDT were pooled and dialysed with 2 mg/ml Tobacco Etch Virus (TEV) protease (prepared in house)[3] in buffer (50 mM MES, 250 mM NaCl, pH 6.5) overnight at 4°C. The dialysed mixture was loaded onto a 5 mL HisTrap column and the flow-through fractions were collected and analysed by SDS-PAGE. Fractions containing purified *Ct*NDT were pooled, flash frozen and stored at -80 °C (Figure S1). Single mutants *Ct*$NDT_{A9S}$, *Ct*$NDT_{V85S}$, *Ct*$NDT_{Y7A}$, *Ct*$NDT_{P40A}$, double mutant *Ct*$NDT_{Y7F\text{-}A9S}$ and triple mutants *Ct*NDT$_{Y7F-A9S-V85}$ and *Ct*NDT$_{Y7F-A9S-P40A}$ were expressed and purified using the same method. Protein concentration was determined by the extinction coefficient at 280nm calculated using Expasy protparam tool[4].

Fractions containing pure *Ct*NDT and *Bp*NDT mutants (>95%) were pooled, flash-frozen and used in the subsequent experiments. Figure S7 depicts all the intact masses for the proteins purified here.

**Substrate specificity of unusual nucleosides and bases using *Ct*NDT$_{Y7F-A9S}$ detected by LC-MS.**

A standard sample contained CtNDT$_{Y7F-A9S}$ (1 µM), 1 mM nucleoside substrate and 1 mM base substrate (details in Table S3) in a mixed solution (50 µL total) (30 mM CHES, MES and HEPES, pH 6.5) incubated at 65 °C for 2 hours. The reaction was quenched with 200 µL of MeOH and incubated at – 80 °C for 15 minutes. Samples were centrifuged at 24000 x g for 10 minutes, and 200 µL of supernatant was removed and evaporated under a stream of nitrogen gas. Sample were resuspended with 75 µL of LC-MS grade $H_2O$ and 10 µL was injected and analysed by LC-MS.

LC conditions: Nucleosides were analysed on a Waters Acquity HSS T3, 1.8 µm, 2.1mm X 100 mm column. A flow rate of 300 µL min$^{-1}$ at initial conditions 99% $H_2$ O + 0.1% Formic Acid with 1% ACN + 0.1% Formic Acid until 1 min, followed by a gradient step to 50% $H_2$ O + 0.1% Formic Acid with 50% ACN + 0.1% Formic Acid to 3.5 mins. A cleaning step was carried out with 1% $H_2O$ + 0.1% Formic Acid with 99% ACN + 0.1% Formic Acid for 5.5 mins and then was re-equilibrated to 99% $H_2O$ + 0.1% Formic Acid with 1% ACN + 0.1% Formic Acid to 7 mins. The column temperature was 40°C throughout the run.

MS conditions: Samples were analysed using a Waters Q-ToF Xevo G2XS, using MS$^E$ for data acquisition. The capillary voltage was set at 2.5 kV in positive ion mode. The source and desolvation gas temperatures of the mass spectrometer were set at 120 °C and 500 °C, respectively. The cone gas flow was set to 50 L/hr, whilst the desolvation gas flow was set at 1000 L/hr. An MS$^E$ scan was performed between 50 – 700 *m/z* where function 1 employed

MS analysis whilst function 2 applied a collision energy ramp from 15 to 30 V to perform MS/MS fragmentation. A lock spray signal was measured, and a mass correction was automatically applied by collecting every 10s, averaging 3 scans of 1s each using Leucine Enkephalin as a standard (556.2771 *m/z*).

**Standard enzyme kinetics of CtNDT with nucleoside substrates using HPLC endpoint assay**

Deoxynucleosides assay: A standard assay contained CtNDT (10 nM) or $CtNDT_{Y7F}$ (50 nM), 10 µM-1000 µM dGuo and a nucleobase (10 mM Ade) in a mixed solution (50 µL total) (30 mM CHES, MES and HEPES, pH 6.5) incubated at 45 °C for 5, 10 and 15 minutes. Michaelis-Menten curves for UricA contained 2 µM CtNDT, 1000 µM 2′d-Ado, UricA (varying as follows: 10, 25, 50, 100, 250, 500, 1000, 2500 µM) in 30 mM CHES, MES and HEPES, pH 6.5.

Ribonucleosides assay: A standard assay contained enzyme [different concentrations of each variant used as follows: CtNDT (1 µM), $CtNDT_{Y7F}$ (0.5 µM for RGua, 1 µM for RIno, 0.8 µM for RAdo), $CtNDT_{Y7A}$ (0.8 µM), *Ct*$NDT_{Y7F-A9S}$ (1 µM), triple mutants *Ct*$NDT_{Y7F-A9S-V85S}$ and *Ct*$NDT_{Y7F-A9S-P40A}$ (both at 1 µM), $CtNDT_{P40A}$ (1 µM), $CtNDT_{D62N}$ (1 µM)], varying 10 µM-1000 µM ribonucleoside (RAdo, RGua or RIno) and a base (10 mM Hyp or Ade) in a mixed solution (50 µL total) (30 mM CHES, MES and HEPES, pH 7). Reactions were incubated at 65 °C for 5, 10 and 15 minutes. At different time points, the reaction mixture was quenched with 200 µL of 10 M Urea and centrifuged at 24000 x g for 10 minutes. 100 µL of the mixture was placed into a 96-well round bottom microplate (Agilent Technologies) and 10 µL of each sample was injected into the column in HPLC (Shimadzu) or UHPLC (Thermo Fisher Scientific). All the experiments were done in duplicate. The data were analysed with Chromeleon ver 6.80 or LabSolution ver 5.11 and were fitted and plotted as Michaelis-Menten Equation using Prism 9.0.

HPLC conditions: A HSS T3, 2.5 µm, 50 mm X 4.6 mm column (Waters™) was used with buffer A (10 mM trimethyl ammonium acetate, pH 7) and buffer B (ACN + 0.1% TFA). A gradient elution was used to separate the compounds: at 0-10 mins, 99% to 85% buffer A and 1% to 15% buffer B, and 10-15 mins,

85% to 0% buffer A and 15% to 100% buffer B. The oven temperature was set at 40 °C and the absorbance was set at 260 nm. The column was equilibrated with buffer A : buffer B (1:99) for at least 15 mins before each injection. Retention times for the reference natural compounds: 2'- dAdo: 8.2 min; 2'-dGuo: 3.8 min; 2'- dIno: 2.5 min; RAdo: 8.1 min; RGuo: 2.5 min; RIno: 2.8 min; Ade: 2.4 min; Gua: 1.4 min; Hyp: 1.4 min.

#### Estimation of reaction yield by HPLC

Compound identities were confirmed by comparison with commercial standards for both the nucleoside products and the starting materials. Quantification was performed using calibration curves generated with authentic standards. Chromatographic analysis was conducted as previously described, with detection at 260 nm. Peak areas for reactants and products were integrated, and UV absorbance values were converted to molar quantities using the appropriate calibration curves for each standard.

#### Determination of $K_D$ for Immucillin-H by differencial scanning fluorimetry

Assays were prepared in a MicroAmp™ 96-well optical reaction plate (Fisher Scientific) with a final volume of 20 µL per well, containing buffer (30 mM MES, 250 mM NaCl, pH 6.5) and 5X SYPRO Orange dye (Invitrogen). The concentration of Immucillin-H was varied in a serial dilution, the test plate was sealed with adhesive film and centrifuged for 5 minutes at 500 rpm. Fluorescence emission was recorded using an Applied Biosystems® QuantStudio 1 Real-Time PCR system (λex = 520 nm, λem = 558 nm). Temperature increases were performed over the range of 25–95 °C, with increments of 1 °C/min, individual metling curves were analysed using Applied Biosystems Protein Thermal Shift Software v.1.4 (Thermo Fisher Scientific) to determine individual $T_M$ values. The $K_D$ was determined in GraphPad Prism by using the following equation[5].

$$Y = Bottom + (Top - Bottom)(1 - \frac{P - K_D - X + \sqrt{(P + X + K_D)^2 - 4PX}}{2P})$$

(P: protein concentration; $K_D$: dissociation constant; Top: maximum temperature; Bottom: minimum temperature; X: ligand concentration)

**Crystallisation, data collection, structure solution, model building, refinement and validation**

CtNDT$_{Y7F}$ with Immucillin-H – The protein concentration was 10 mg/mL, and ligand Immucillin-H was used at a final concentration of 1 mM. Co-crystals were grown at 20 °C using the sitting drop vapour diffusion technique, with a drop size of 0.3 µl in a 1:1, 1:2 or 2:1 reservoir: protein solution. Crystallisation conditions were from the commercial screen PACT (well D12): 20% w/v PEG 6000; 0.1 M Tris pH 8.0; 0.002 M zinc chloride. Crystals were cryoprotected in mother liquor supplemented with 20% (v/v) ethylene glycol before being flash cooled in liquid nitrogen. Diffraction data were collected at the Diamond Light Source in Oxford, UK, on I04 beamlines. Data reduction and processing were completed using the xia2 suite.

CtNDT$_{Y7F_A9S}$ with cordycepin – The protein concentration was 10 mg/mL, and ligand cordycepin was used at a final concentration of 1 mM. Co-crystals were grown at 20 °C using the sitting drop vapour diffusion technique, with a drop size of 0.3 µl in a 1:1, 1:2 or 2:1 reservoir: protein solution. Crystallisation conditions were 21.429% PEG Smear, 0.1 M Bis-Tris Propane pH 8.33. Crystals were cryoprotected in mother liquor supplemented with 20% (v/v) ethylene glycol before being flash cooled in liquid nitrogen. Diffraction data were collected at the Diamond Light Source in Oxford, UK, on I04 beamline. Data reduction and processing were completed using the xia2 suite. The structure was solved by molecular replacement with PHASER searching for four monomers in the ASU. The search component was CtNDT (pdb 8PQS), which had been modified sculptor and manually truncated in COOT.

For both, structures were solved by molecular replacement with PHASER searching for four monomers in the ASU. The search component was CtNDT (pdb 8PQS), which had been modified sculptor and manually truncated in COOT. Protein structures were built/ modified using COOT, with cycles of refinement in PHENIX. Restraints for the ligand were generated using

ACEDRG, available at the CCP4 cloud server. Crystallographic data are shown in Table S4.

#### Supporting Tables

##### Table S1. Primers used for mutagenesis

| Name | 5’-3’ Primer Sequence |
|---|---|
| $CtNDT_{Y7F}$ Forward | TATTTTCCTTGCCAGCCCTTATGGTTTTAG |
| $CtNDT_{Y7F}$ Reverse | CAAGGAAAATAATCTTACGTTTCATCCCCTGG |
| $CtNDT_{Y7A}$ Forward | TAAGATTATTGCCCTTGCCAGCC |
| $CtNDT_{Y7A}$ Reverse | CGTTTCATCATTCCTGTAAG |
| $CtNDT_{P40A}$ Forward | GGTTTGGGAAGCATTTGCTCGTAATAAC |
| $CtNDT_{P40A}$ Reverse | TCGATCCCCAGTGCCTCT |
| $CtNDT_{Y7F-A9S}$ Forward | TATTTTCCTTAGCAGCCCTTATGG |
| $CtNDT_{Y7F-A9S}$ Reverse | ATCTTACGTTTCATCATTCC |
| $CtNDT_{Y7F-A9S-V85S}$ Forward | AGATGAGGGCAGTATGGTAGAATTGGGTATGGC |
| $CtNDT_{Y7F-A9S-V85S}$ Reverse | GGGGGCGTTCCATTAACC |
| BpNDT NDT Forward | GAAATAATTTTGTTTAACTTTTGGAATTCCATATG CACCAC |
| BpNDT NDT Reverse | CAAGGGGTTATGCTAGGGGGCCCCAACTTTCA CTTCAC |
| pJ414 NDT Forward | CCCCCTAGCATAACCCCTTG |
| pJ414 NDT Reverse | AAAAGTTAAACAAAATTATTTCTAGAGGGGAATTG |
| $BpNDT_{Y5F}$ Forward | GGCAAAAATCTTTTTGGCGTCAC |
| $BpNDT_{Y5F}$ Reverse | ATTCACTTCACAGCTTTAAG |

**Table S2. Masses expected and observed for nucleosides substrates and products. Only reactions that yielded expected products are shown.**

| Product expected | Nucleoside donor | Nucleobase | Theoretical Mass Nucleoside product [M+H]/[M-H] | Experimental Mass Nucleoside product [M+H]/[M-H] | ppm deviation | Nucleoside product |
|---|---|---|---|---|---|---|
| | | | **WT-*Ct*NDT** | | | |
| 2'-deoxy-2 | 2′-deoxy adenosine | Purine | 237.0982 | 237.0980 | -0.843 | 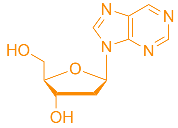 |
| 2'-deoxy-3 | | 2-fluoroadenine | 270.0998 | 270.100 | 0.740 | 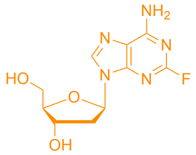 |
| 2'-deoxy-4 | | 6-amino-7H-purin-2-ol (isoguanine) | 268.1051 | 268.105 | -0.373 | 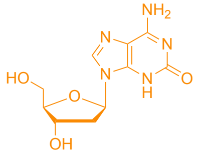 |

| | | | | | | |
|---|---|---|---|---|---|---|
| 2’-deoxy-5 | | 1-methyl-3,7-dihydro-purine-2,6-dione | 283.1038 | 283.104 | 0.706 | 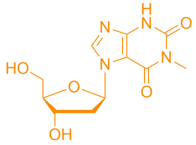 |
| 2’-deoxy-6 | | N-(7H-Purin-6-yl)benzamide | 356.1353 | 356.135 | -0.842 | 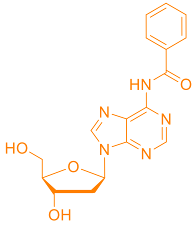 |
| 2’-deoxy-7 | | N, N-dimethyl-7H-purine-6-amine | 280.1408 | 280.141 | 0.714 | 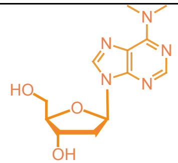 |
| 2’-deoxy-8 | | 5-Nitrobarbituric acid | 290.0618 | 290.062 | 0.690 | 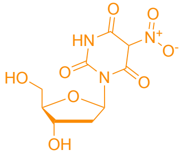 |
| 2’-deoxy-9 | | 6-(benzyloxy)-9H-purine | 343.1408 | 343.140 | -2.331 | 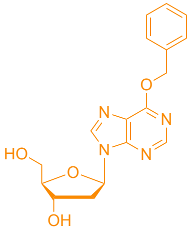 |

| | | | | | | |
|---|---|---|---|---|---|---|
| 2’-deoxy-11 | | 4,5,6,7-tetrahydro-1H-pyrazolo[4,3,-c]pyridine | 240.1348 | 240.1349 | 0.416 | 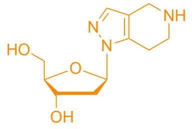 |
| 2’-deoxy-12 | 2′-deoxyguanosine | 2,5,6,7,8,9-hexahydro-3H-cyclohepta[c]pyridazine-3-one | 281.1501 | 281.150 | -0.356 | 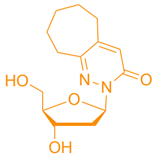 |
| 2’-deoxy-13 | | 1,4,5,6-Tetrahydropyrrolo[3,4-c]pyrazole | 226.1191 | 226.120 | 3.980 | 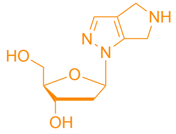 |
| 2’-deoxy-14 | | 5-chlorobenzotriazole | 270.0639 | 270.064 | 0.111 | 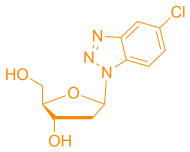 |
| 2’-deoxy-15 | 2′-deoxy adenosine | 6-mercaptopurine | 269.0708 | 269.070 | -2.973 | 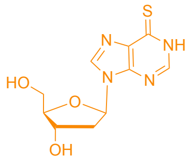 |
| 2’-deoxy-16 | | uric acid* | 283.0679 | 283.068 | 0.353 | 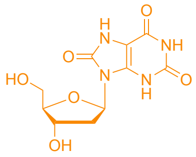 |

| | | | | | | |
|---|---|---|---|---|---|---|
| 2’-deoxy-19 | | 6-chloro-3-methyluracil* | 275.0435 | 275.043 | -1.817 | 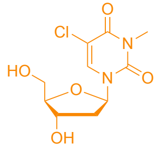 |
| 2’-deoxy-21 | 2′-deoxyuridine | 5-azacytosine | 229.0938 | 229.094 | 0.873 | 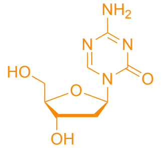 |
| 2’-deoxy-22 | 2′-deoxy adenosine | 3-aminopyridin-2(1H)-one | 227.1026 | 227.102 | -2.677 | 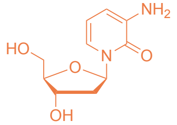 |
| 2’-deoxy-38 | | IBMX | 339.1668 | 339.167 | 0.589 | 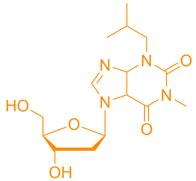 |
| 2’-deoxy-47 | | 4-hydroxy-7-azaindole | 251.1032 | 251.1038 | 2.389 | 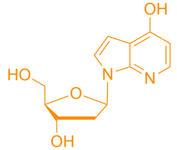 |
| 2’-deoxy-48 | | 5-chloro-7H-pyrrolo[2,3-d]pyrimidine | 270.0645 | 270.0657 | 4.443 | 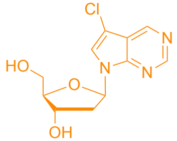 |

| | | | | | | |
|---|---|---|---|---|---|---|
| 2’-deoxy-54 | | 2,6-diaminopurine | 267.1208 | 267.1214 | 2.246 | 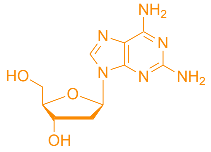 |
| Adenosine | Guanosine | adenine | 268.1040 | 268.105 | 3.711 | 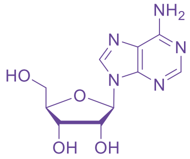 |
| Adenosine | Inosine | adenine | 268.1040 | 268.105 | 3.711 | |
| Cytidine | Adenosine | cytosine | 244.0927 | 244.093 | 0.934 | 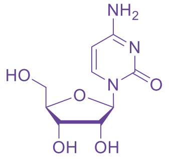 |
| 2'-amino-2’-deoxy-adenosine | 2′-amino-2’-deoxyuridine | adenine | 267.1205 | 267.120 | -1.872 | 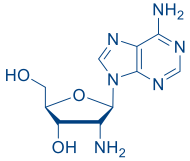 |
| 3′-deoxyguanosine | 3′-deoxyadenosine (Cordycepin) | guanine | 268.104005 | 268.104 | -0.018 | 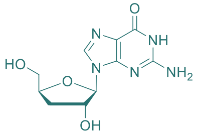 |
| **$CtNDT_{Y7F}$** | | | | | | |

| | | | | | | |
|---|---|---|---|---|---|---|
| Cytidine | Adenosine | cytosine | 244.0927 | 244.093 | 0.934 | 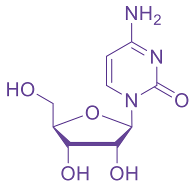 |
| Adenosine | Guanosine | adenine | 268.1040 | 268.105 | 3.711 | 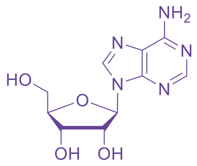 |
| Adenosine | Inosine | adenine | 268.1040 | 268.105 | 3.711 | |
| **CtNDT$_{Y7F-A9S}$** | | | | | | |
| 3’-deoxy-2 | 3′-deoxyadenosine (Cordycepin) | purine | 237.0981 | 237.098 | -0.422 | 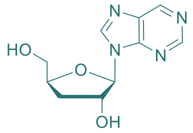 |
| 3’-deoxy-3 | | 2-fluoroadenine | 270.0998 | 270.100 | 0.740 | 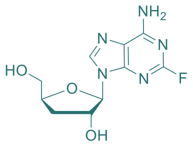 |
| 3’-deoxy-5 | | 1-methyl-3,7-dihydro-purine-2,6-dione | 283.1038 | 283.104 | 0.706 | 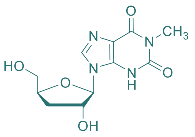 |

| | | | | | | |
|---|---|---|---|---|---|---|
| 3'-deoxy-7 | | N, N-dimethyl-7H-purine-6-amine | 280.1408 | 280.141 | 0.714 | 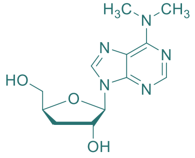 |
| 3'-deoxy-9 | | 6-(benzyloxy)-9H-purine | 343.1408 | 343.141 | 0.582 | 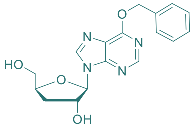 |
| 3'-deoxy-10 | | 1H-Indazole | 235.1077 | 235.108 | 1.301 | 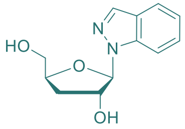 |
| 3'-deoxy-15 | | 6-mercaptopurine | 269.0708 | 269.070 | -2.973 | 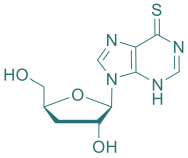 |
| 3'-deoxy-38 | | IBMX | 339.1668 | 339.166 | -2.359 | 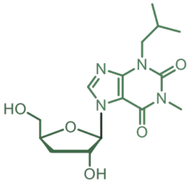 |
| 3'-deoxy-54 | | 2,6-diaminopurine | 267.1208 | 267.1211 | 1.123 | 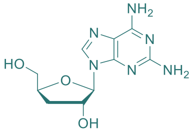 |

| | | | | | | |
|---|---|---|---|---|---|---|
| Ribo-1 | Adenosine | 2,6-dichloropurine | 321.0158 | 321.015 | -2.492 | 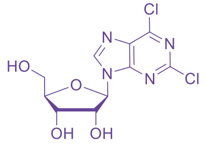 |
| Ribo-2 | | purine | 253.0938 | 253.093 | -3.161 | 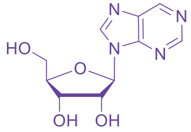 |
| Ribo-3 | | 2-fluoroadenine | 286.0948 | 286.095 | 0.699 | 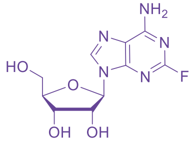 |
| Ribo-5 | | 1-methyl-3,7-dihydro-purine-2,6-dione | 299.0988 | 299.099 | 0.669 | 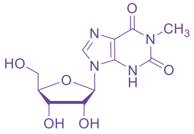 |
| Ribo-6 | | N-(7H-Purin-6-yl)benzamide | 372.1308 | 372.1322 | 3.762 | 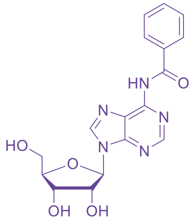 |
| Ribo-7 | | N, N-dimethyl-7H-purine-6-amine | 296.1358 | 296.136 | 0.675 | 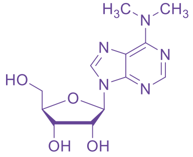 |

| | | | | | | |
|---|---|---|---|---|---|---|
| Ribo-9 | | 6-(benzyloxy)-9H-purine | 359.1358 | 359.135 | -2.228 | 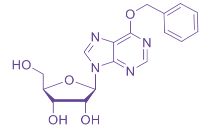 |
| Ribo-10 | | 1H-Indazole | 251.1028 | 251.103 | 0.796 | 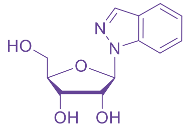 |
| Ribo-13 | | 1,4,5,6-tetrahydropyrrolo[3,4,-c] pyrazole | 242.1138 | 242.1148 | 4.130 | 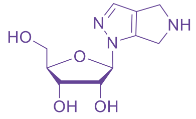 |
| Ribo-14 | | 5-chlorobenzotriazole | 286.0588 | 286.058 | -2.796 | 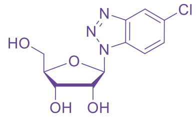 |
| Ribo-15 | | 6-mercaptopurine | 285.0658 | 285.065 | -2.806 | 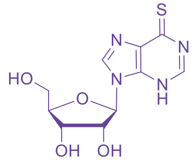 |
| Ribo-38 | | IBMX | 355.1618 | 355.162 | 0.563 | 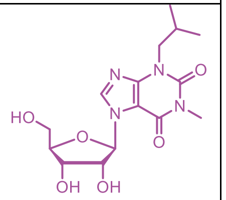 |

| | | | | | | |
|---|---|---|---|---|---|---|
| ribo-47 | | 4-hydroxy-7-azaindole | 267.0978 | 267.0969 | -3.369 | 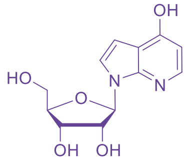 |
| ribo-54 | | 2,6-diaminopurine | 283.1158 | 283.1163 | 1.766 | 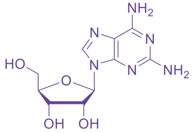 |
| | **WT-*Bp*NDT** | | | | | |
| 2’-deoxy-18 | 2′-deoxyguanosine | 5-fluorouracil* | 245.0572 | 245.057 | -0.816 | 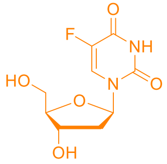 |
| 2’-deoxy-20 | | isocytosine | 228.0988 | 228.098 | -3.507 | 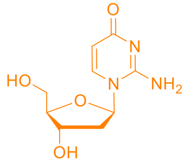 |
| 2’-deoxy-21 | | 5-azacytosine | 229.0938 | 229.094 | 0.873 | 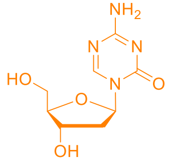 |

| | | | | | | |
|---|---|---|---|---|---|---|
| 2’-deoxy-23 | | 2-thiouracil* | 243.0442 | 243.044 | -0.823 | 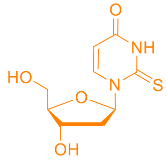 |
| 2’-deoxy-24 | | 6-amino-3-methyluracil* | 256.0933 | 256.094 | 2.733 | 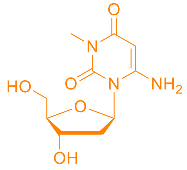 |
| 2’-deoxy-26 | | pyrimidin-4(3H)-one | 213.0878 | 213.087 | -3.754 | 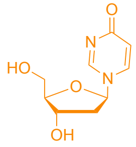 |
| 2’-deoxy-29 | | 5-chlorouracil* | 261.0282 | 261.027 | -4.597 | 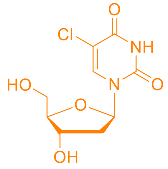 |
| 2’-deoxy-30 | | 5-iodouracil* | 352.9632 | 352.963 | -0.566 | 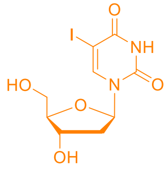 |

| | | | | | | |
|---|---|---|---|---|---|---|
| 2’-deoxy-32 | | 5-bromouracil* | 306.9928 | 306.992 | -2.606 | 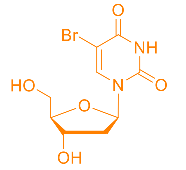 |
| 2’-deoxy-43 | | 5-(2-chloroethyl)-1H-tetrazole* | 247.0592 | 247.059 | -0.810 | 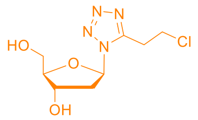 |
| | **BpNDT Y5A** | | | | | |
| Ribo-18 | Guanosine | 5-fluorouracil | 261.0522 | 261.052 | -0.766 | 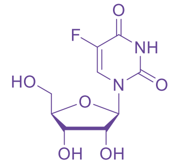 |

*negative mode

**Table S3. Steady-state parameters for enzyme variants** with 2′-deoxyribonucleosides and ribonucleosides.

Concentrations of each variant used as follows: CtNDT (1 µM), CtNDT$_{Y7F}$ (0.5 µM for RGua, 1 µM for RIno, 0.8 µM for RAdo), CtNDT$_{Y7A}$ (0.8 µM), *Ct*NDT$_{Y7F\text{-}A9S}$ (1 µM), triple mutants *Ct*NDT$_{Y7F\text{-}A9S\text{-}V85S}$ and *Ct*NDT$_{Y7F\text{-}A9S\text{-}P40A}$ (both at 1 µM), CtNDT$_{P40A}$ (1 µM), CtNDT$_{D62N}$ (1 µM).

| **Enzyme** | **Donor** | **Acceptor** | $K_M$ **(mM)** | $k_{cat}$ **(s$^{-1}$)** | $k_{cat}/K_M$ **(mM$^{-1}$s$^{-1}$)** |
|---|---|---|---|---|---|
| CtNDT | 2′-dGuo | Ade | 0.69 ± 0.14 | 6.91 ± 0.72 | 10.01 ± 2.28 |
| | RAdo | Hyp | 0.25 ± 0.05 | 0.03 ± 0.02 | 0.12 ± 0.09 |
| | RGua | Ade | 1.09 ± 0.31 | 0.06 ± 0.01 | 0.06 ± 0.02 |
| | RIno | | 0.92 ± 0.24 | 0.01 ± 0.01 | 0.01 ± 0.01 |
| | 2′-dAdo | Uric acid | 1.46 ± 0.21 | 0.32 ± 0.02 | 0.22 ± 0.03 |
| CtNDT$_{Y7F}$ | 2′-dGuo | Ade | 0.67 ± 0.08 | 0.21 ± 0.01 | 0.32 ± 0.40 |
| | RAdo | Hyp | 0.11 ± 0.01 | 0.04 ± 0.01 | 0.33 ± 0.03 |
| | RGua | Ade | 0.10 ± 0.02 | 0.02 ± 0.01 | 0.20 ± 0.04 |
| | RIno | | 0.71 ± 0.17 | 0.03 ± 0.01 | 0.04 ± 0.01 |
| CtNDT$_{Y7A}$ | RIno | Ade | 0.53 ± 0.01 | 0.02 ± 0.01 | 0.04 ± 0.01 |
| *Ct*NDT$_{Y7F\text{-}A9S}$ | | | 0.62 ± 0.10 | 0.05 ± 0.01 | 0.08 ± 0.04 |
| *Ct*NDT$_{Y7F\text{-}A9S\text{-}V85S}$ | | | 0.84 ± 0.21 | 0.04 ± 0.01 | 0.05 ± 0.02 |
| *Ct*NDT$_{Y7F\text{-}A9S\text{-}P40A}$ | | | 0.06 ± 0.01 | 0.004 ± 0.001 | 0.07 ± 0.01 |
| CtNDT$_{P40A}$ | | | 0.10 ± 0.02 | 0.01 ± 0.01 | 0.07 ± 0.02 |
| CtNDT$_{D62N}$ | | | 0.80 ± 0.09 | 0.002 ± 0.001 | 0.003 ± 0.001 |

**Table S4: Crystallization conditions**

| PDB accession code | Protein | Protein concentraiton | **Ratio protein/precipitant** | **Crystallization condition** | **Ligand and concentration for co-crystallization** |
|---|---|---|---|---|---|
| 9EMW | Y7F | 10 mg/ml | 1:1 | 0.1 M Tri, pH 8.0, 20% PEG 6000, 0.002 M zinc chloride | 1 mM ImmH, co-crystals |
| 9EMX | DM (Y7F A9S) | 10 mg/ml | 1:1 | 0.1 M Bis-Tris Propane, pH 8.33, 21.429% PEG Smear High, 0.05 M magnesium chloride | 1 mM Cordycepin, co-crystals |

#### Table S5: Crystallographic data

| Property | Value | Value |
|---|---|---|
| **pdb code** | 9EMX - *Ct*NDT$_{Y7F-A9S}$ bound to Cordycepin | 9EMW - CtNDTY7F bound to ImmH |
| Space group | P 32 | P 63 |
| Cell constants | 97.38Å 97.38Å 66.46Å | 135.76Å 135.76Å 87.55Å |
| a, b, c, *α*, *β*, *γ* | 90.00◦ 90.00◦ 120.00◦ | 90.00◦ 90.00◦ 120.00◦ |
| Resolution (Å) | 42.17 – 1.77 | 48.8 = 2.51 |
| | 84.33 – 1.77 | 70.22 - 2.51 |
| % Data completeness | 94.0 (42.17-1.77) | 99.8 (48.80-2.51) |
| (in resolution range) | 99.0 (84.33-1.77) | 99.8 (70.22-2.51) |
| R*merge* | 0.18 | 0.16 |
| < I/σ(I) > 1 | 0.96 (at 1.76Å) | 1.23 (at 2.51Å) |
| Refinement program | PHENIX 1.19.2-4158 | PHENIX (1.21.2-5419) |
| R, R*free* | 0.188 , 0.211 | 0.212, 0.262 |
| | 0.187 , 0.210 | 0.213, 0.26 |
| $R_{free}$ test set | 2064 reflections (3.01%) | 1566 reflections (4.98%) |
| CC1/2 | 1 | 1 |
| Wilson B-factor (Å2) | 34.9 | 61.4 |
| Anisotropy | 0.382 | 0.262 |
| Bulk solvent $k_{sol}$(e/Å3), $B_{sol}$(Å2) | 0.31 , 40.2 | 0.31 , 37.1 |
| L-test for twinning2 | < \|*L*\| > = 0.49, < $L^2$ > = 0.33 | < \|*L*\| > = 0.51, < $L^2$ > = 0.34 |
| Estimated twinning fraction | 0.014 for -h,-k,l | 0.035 for h,-h-k,-l |
| | 0.033 for h,-h-k,-l | |
| | 0.018 for -k,-h,-l | |
| $F_o$,$F_c$ correlation | 0.97 | 0.93 |
| Total number of atoms | 5540 | 5227 |
| All atom clashscore | 8.4 | 5.4 |
| Average B, all atoms (Å2) | 41 | 66 |
| Clashscore all atoms | 5.39 | 4,76 |
| Molprobity score | 1.29 | 1.57 |

#### Supporting Figures

**Figure S1: Data for reactions to produce nucleosides.** Masses indicated in red were assigned to nucleoside products and absent from reactions where bases were omitted. For all graphs “intensity” is in arbitrary ion counts. Left, total ion counts for data acquired in selected ion recording (SIR) mode with m/z set for each expected product with a charge of +1. Right, m/z features contributing to the SIR channel acquired, with the expected m/z shown in red.

##### CtNDT WT

Product 2’-deoxy-2

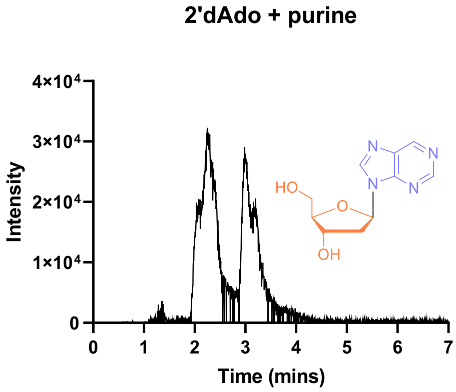

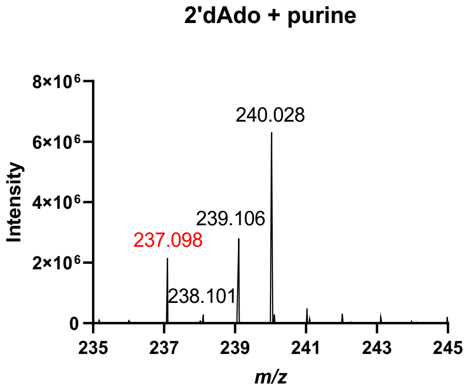

Product 2’-deoxy-3

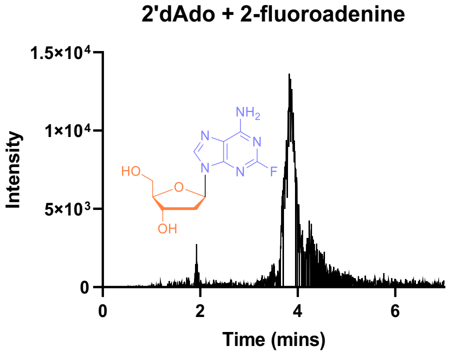

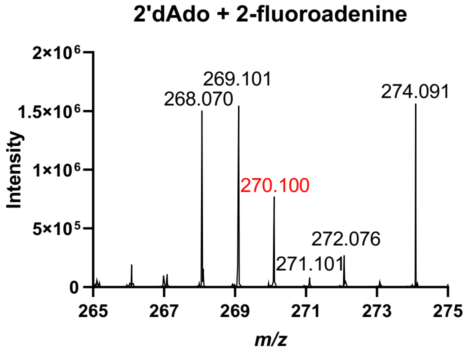

Product 2’-deoxy-4

### Figure S1: Data for reactions to produce nucleosides.

**2'dAdo + isoguanine**

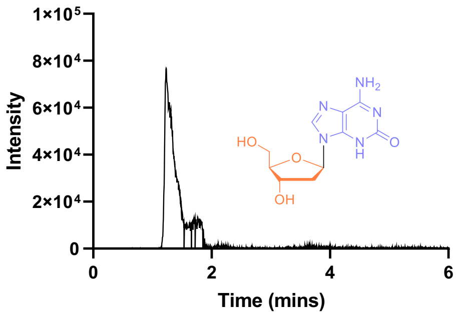

**2'dAdo + isoguanine**

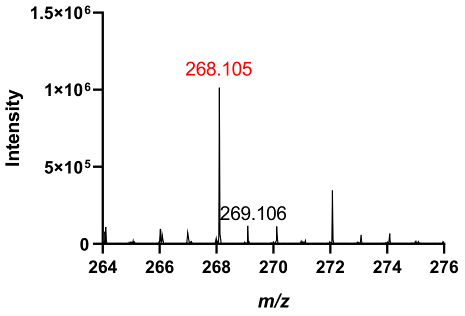

Product 2’-deoxy-5

**2'dAdo + 1-methyl-3,7-dihydro-purine-2,6-dione**

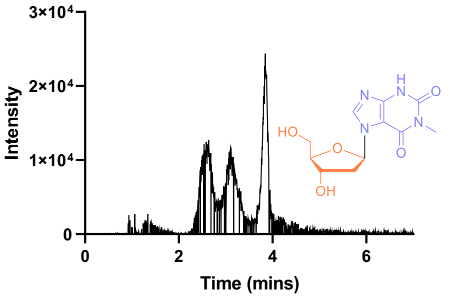

**2'dAdo + 1-methyl-3,7-dihydro-purine-2,6-dione**

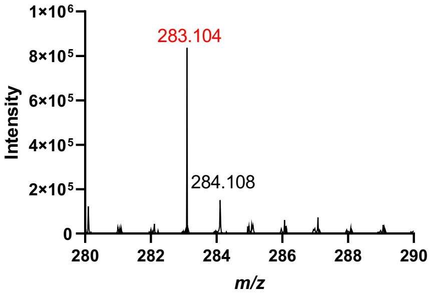

Product 2’-deoxy-6

**2'dAdo + N-(7H-Purin-6-yl)benzamide**

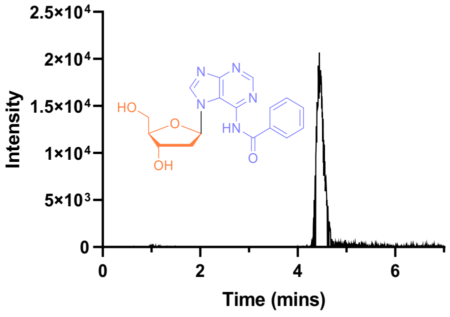

**2'dAdo + N-(7H-Purin-6-yl)benzamide**

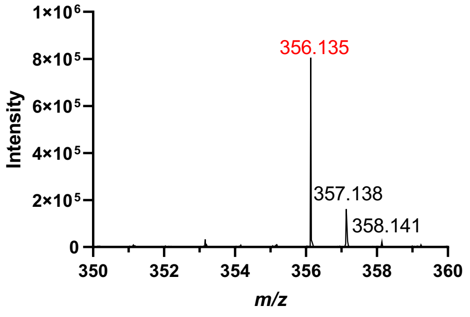

Product 2’-deoxy-7

**2'dAdo + N,N-dimethyl-7H-purine-6-amine**

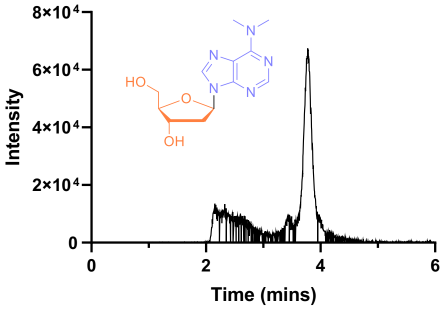

**2'dAdo + N,N-dimethyl-7H-purine-6-amine**

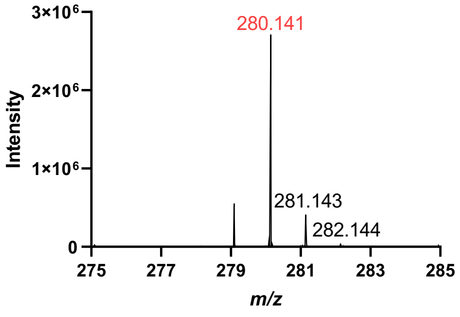

Product 2’-deoxy-8

#### Figure S1: Data for reactions to produce nucleosides.

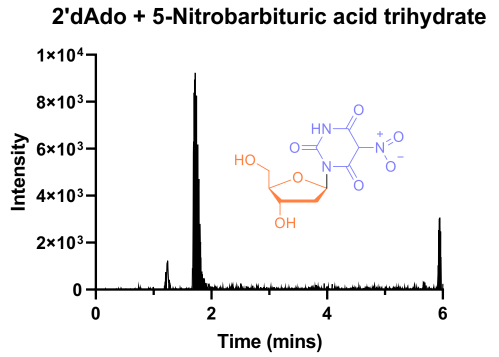

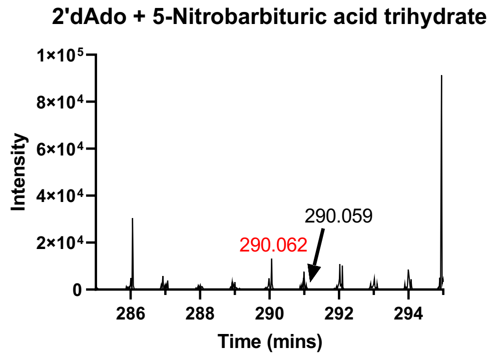

Product 2’-deoxy-9

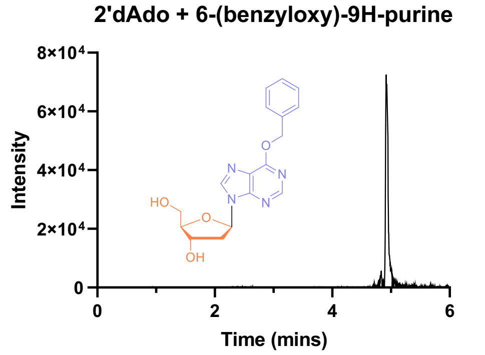

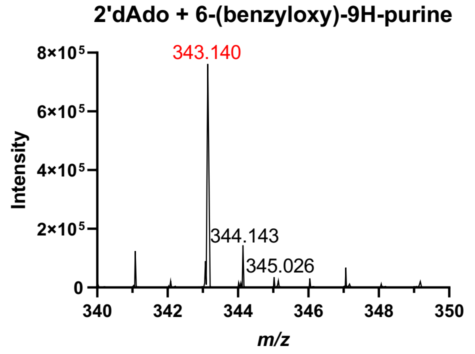

Product 2’-deoxy-11

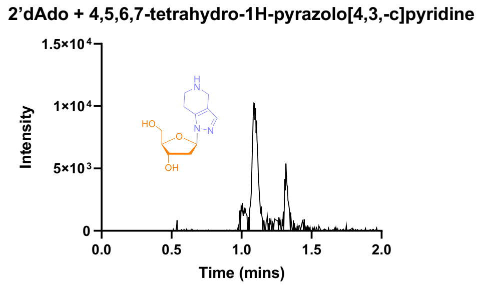

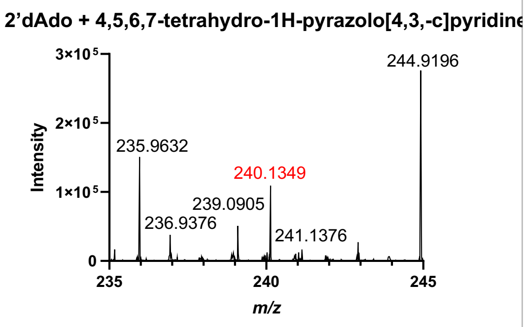

Product 2’-deoxy-12

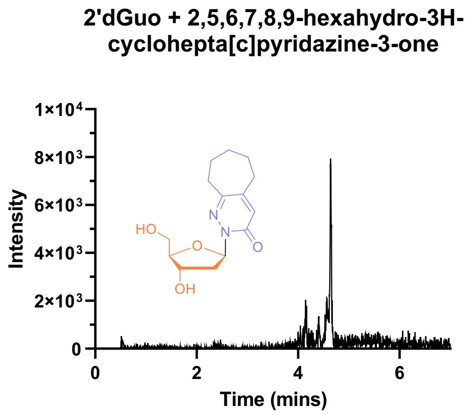

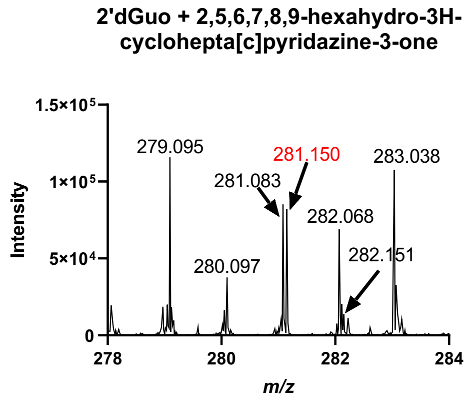

#### Figure S1: Data for reactions to produce nucleosides.

Product 2’-deoxy-13

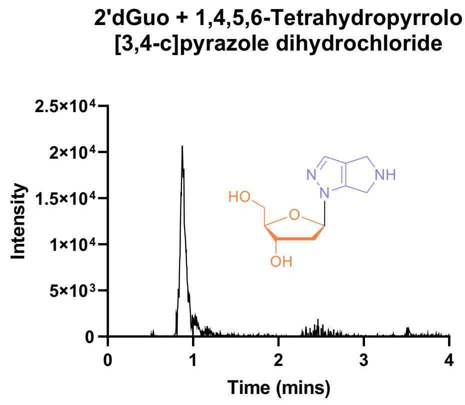

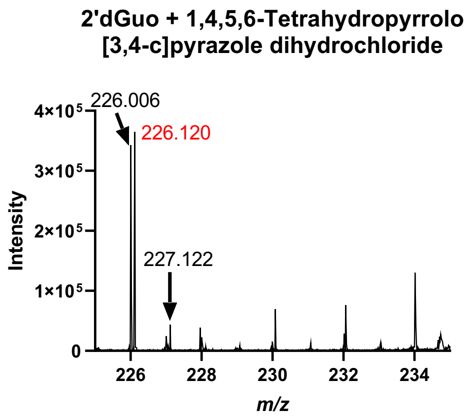

Product 2’-deoxy-14

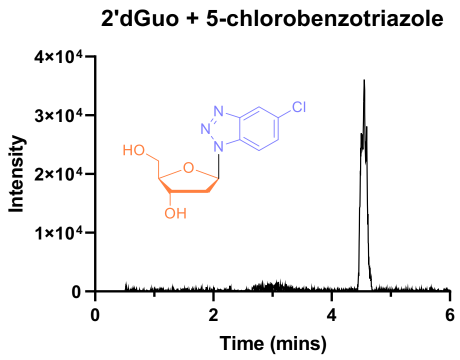

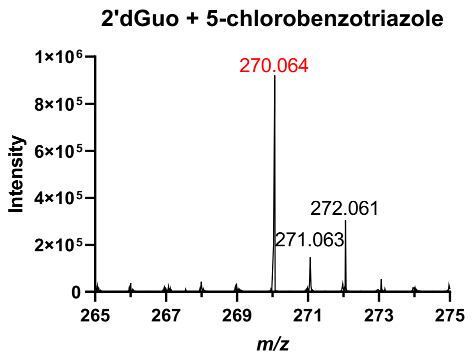

Product 2’-deoxy-15

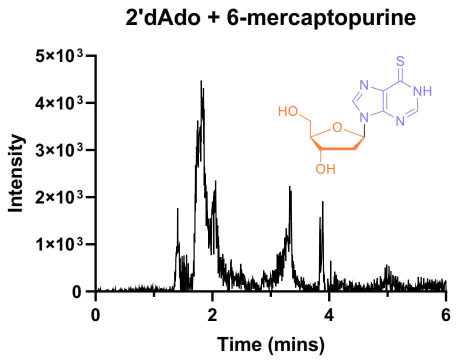

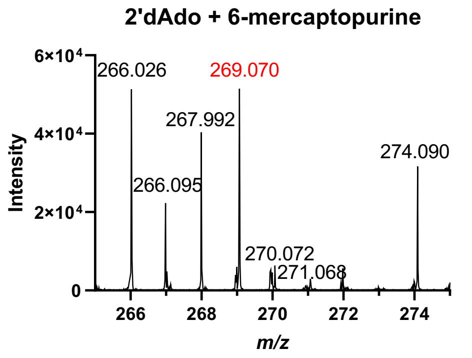

Product 2’-deoxy-16

#### Figure S1: Data for reactions to produce nucleosides.

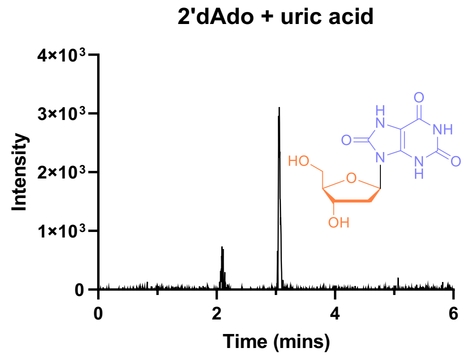

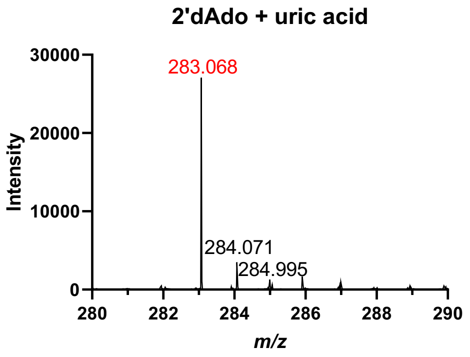

Product 2’-deoxy-19

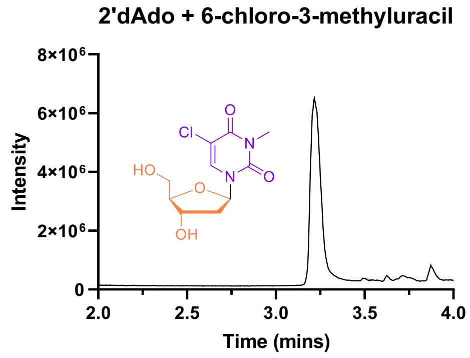

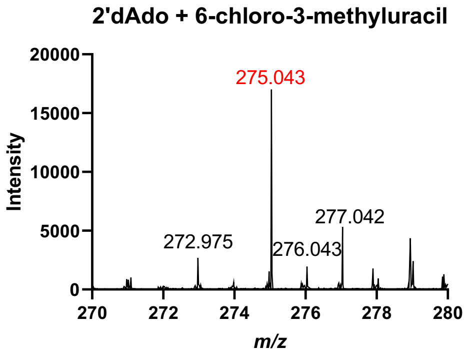

Product 2’-deoxy-21

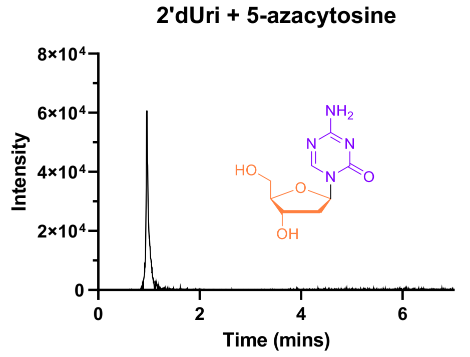

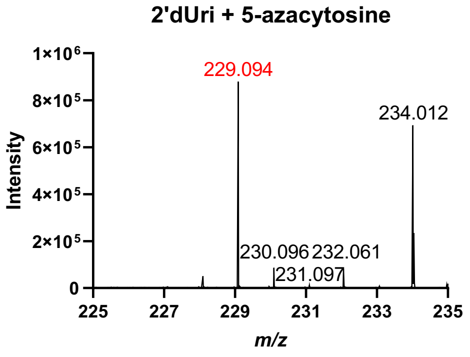

Product 2’-deoxy-22

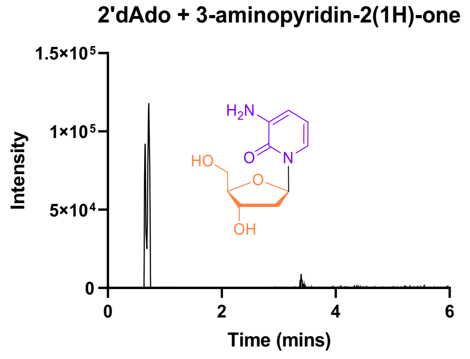

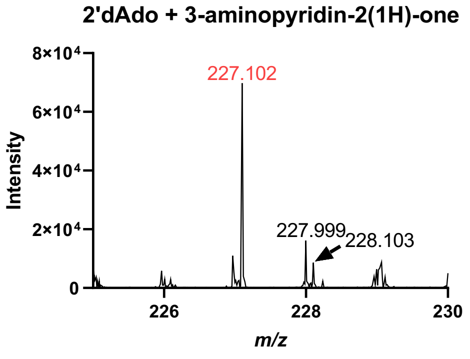

Product 2’-deoxy-38

#### Figure S1: Data for reactions to produce nucleosides.

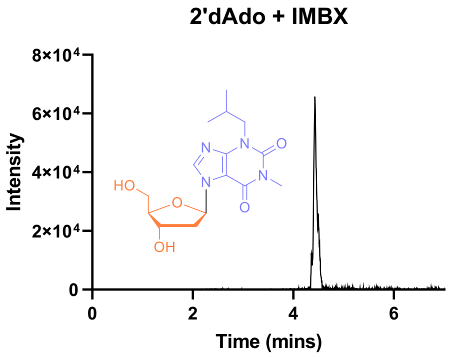

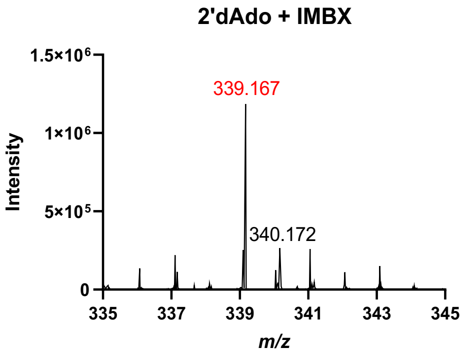

Product 2’-deoxy-47

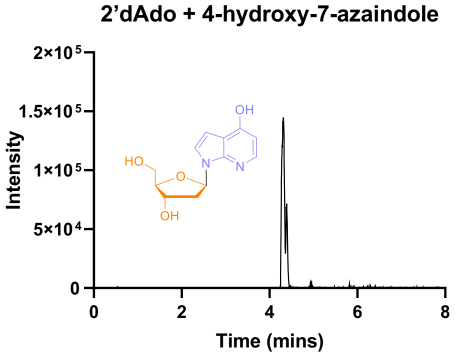

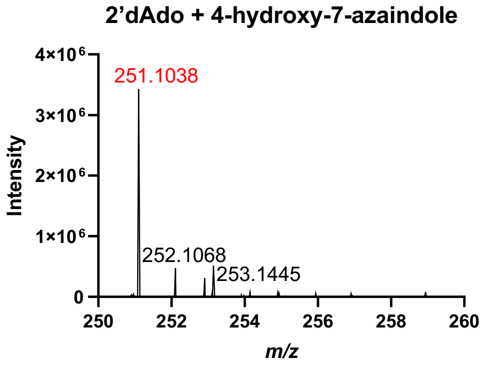

Product 2’-deoxy-48

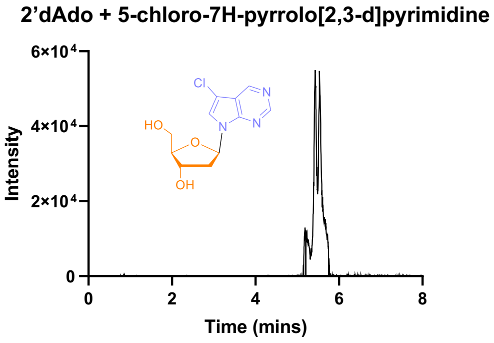

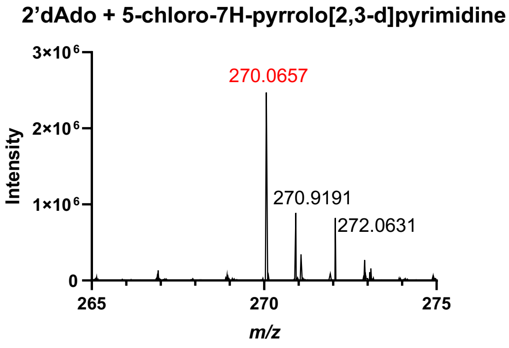

Product 2’-deoxy-54

**Figure S1: Data for reactions to produce nucleosides.**

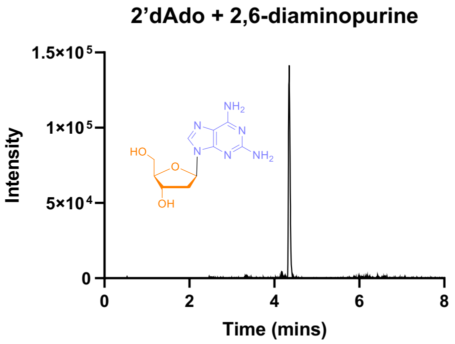

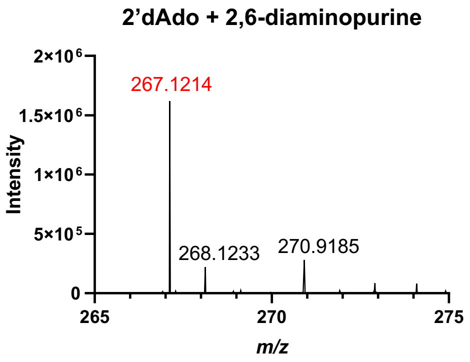

Standard nucleobase substrates

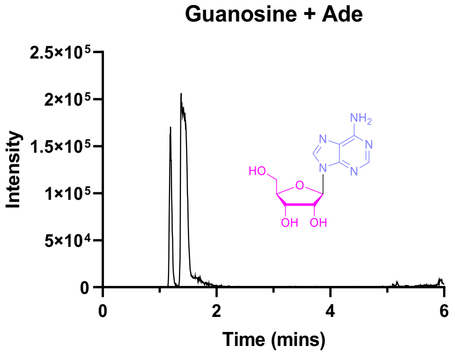

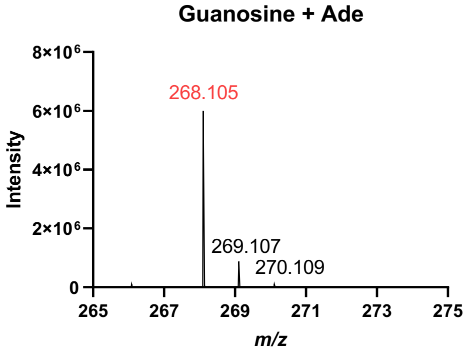

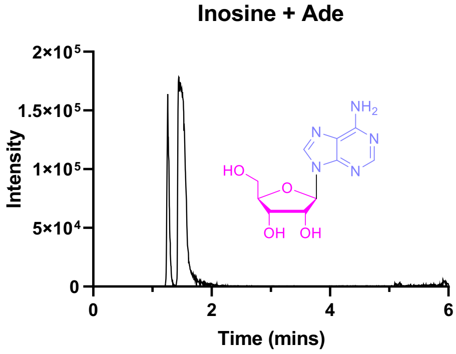

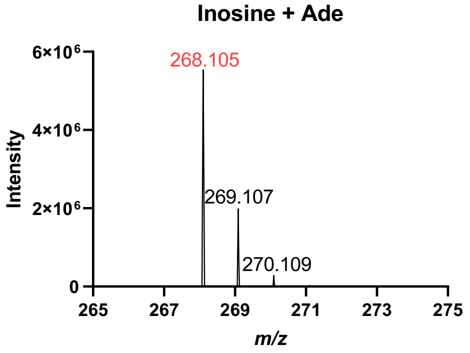

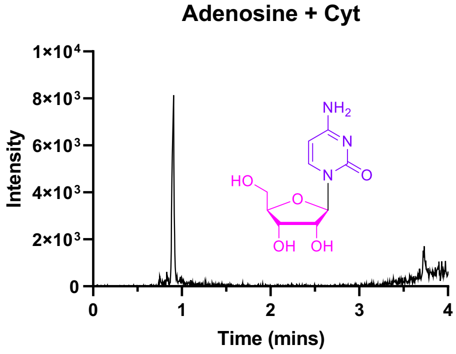

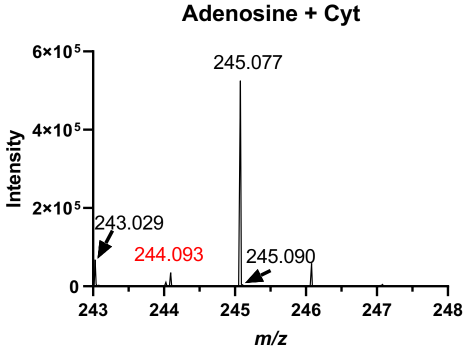

2'-amino-2’-deoxy-adenosine

#### Figure S1: Data for reactions to produce nucleosides.

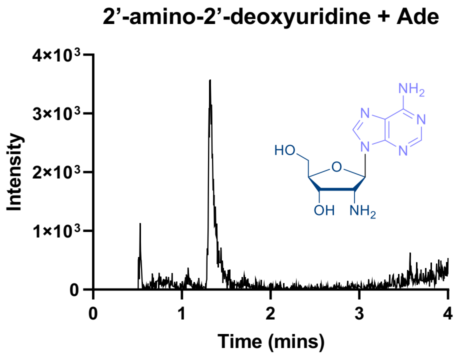

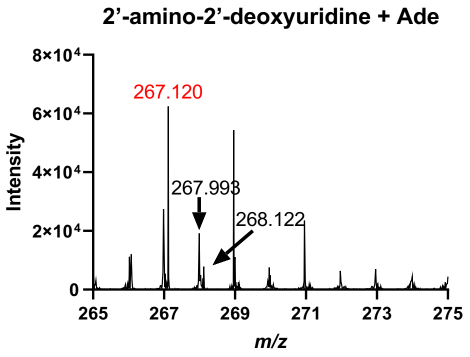

3’-deoxy-adenosine (cordycepin)

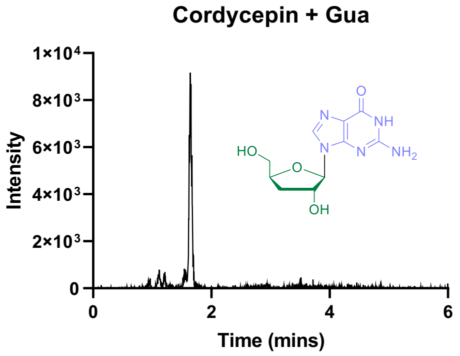

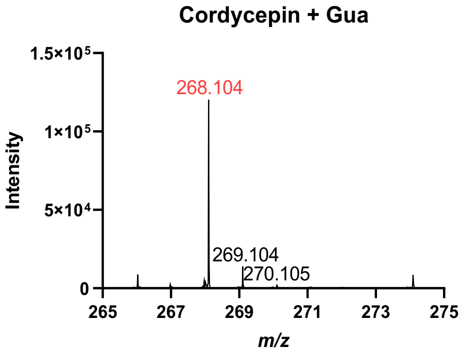

#### CtNDT Y7F – confirmation of activity with ribonucleosides

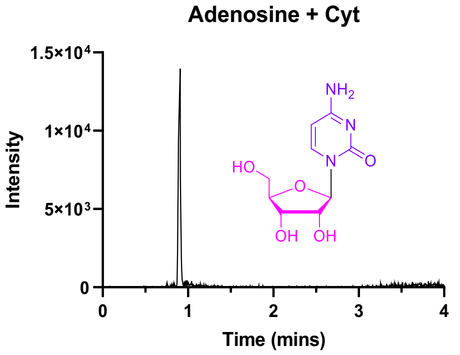

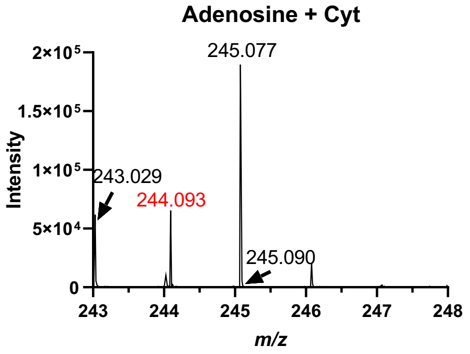

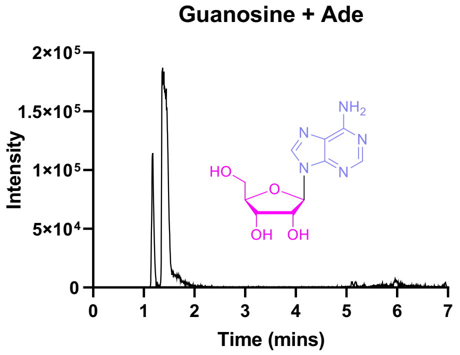

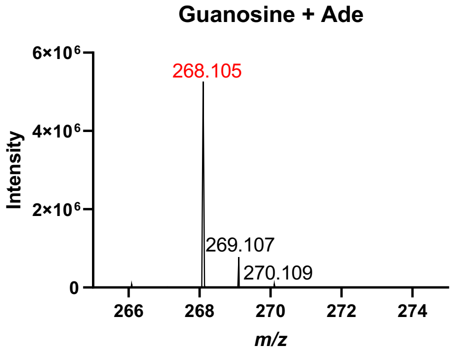

#### Figure S1: Data for reactions to produce nucleosides.

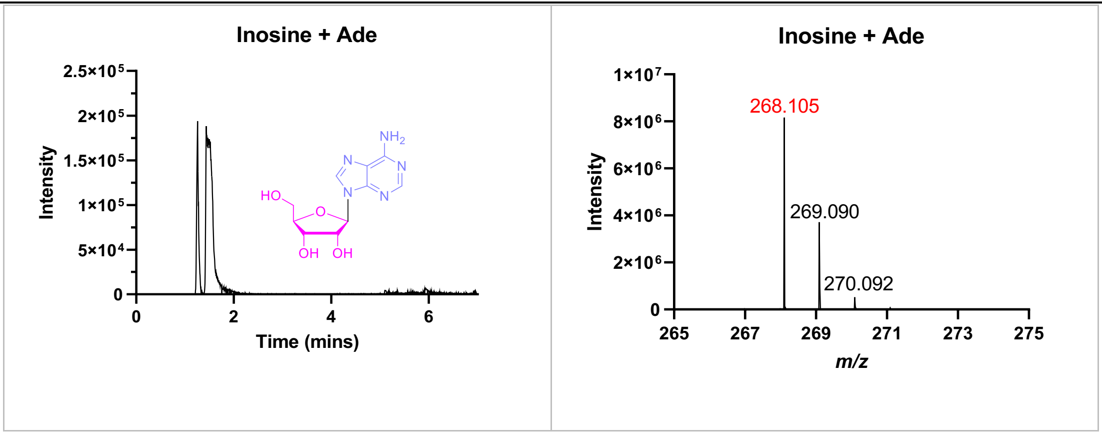

#### Figure S1: Data for reactions to produce nucleosides.

##### CtNDT$_{Y7F\text{-}A9S}$

Product 3’-deoxy-2

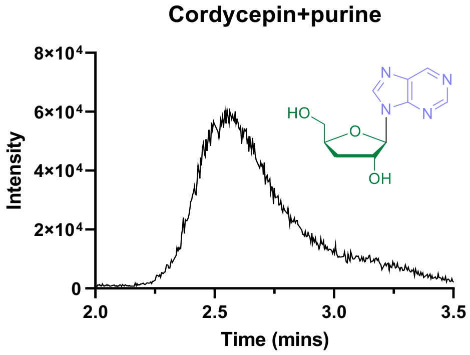

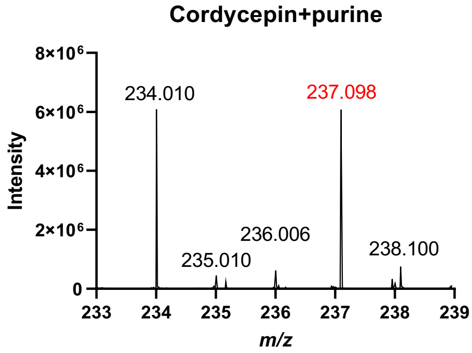

Product 3’-deoxy-3

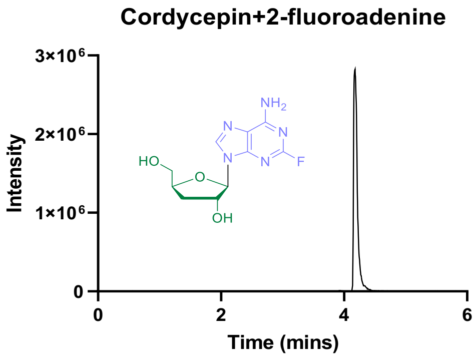

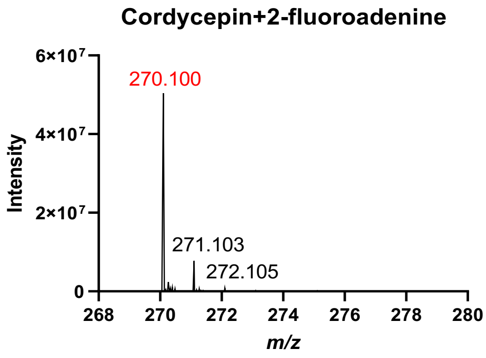

Product 3’-deoxy-5

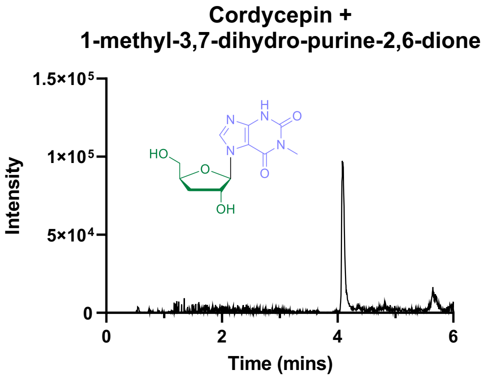

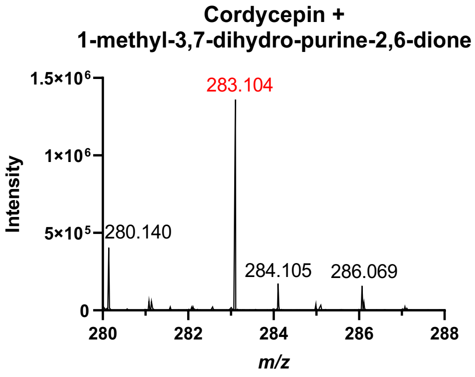

#### Figure S1: Data for reactions to produce nucleosides.

Product 3'-deoxy-7

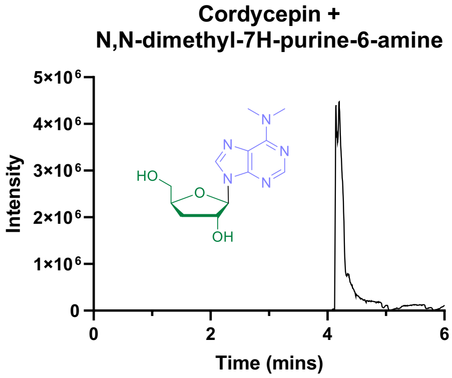

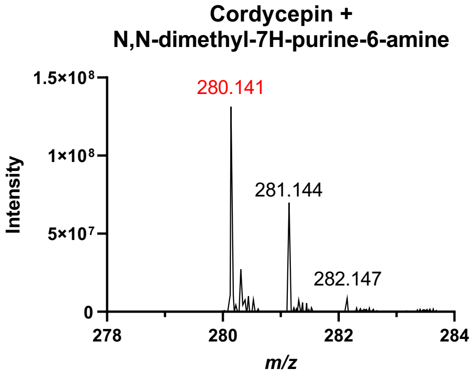

Product 3'-deoxy-9

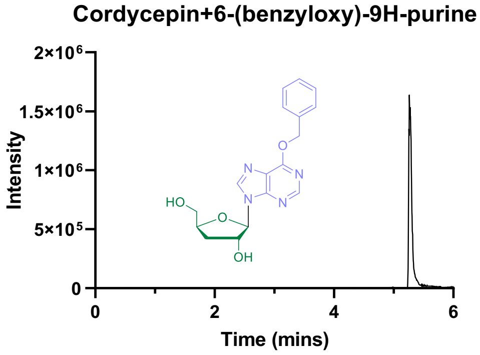

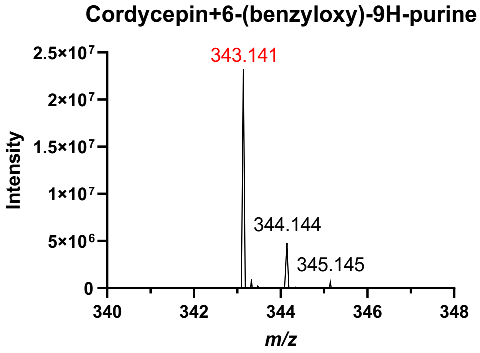

Product 3'-deoxy-10

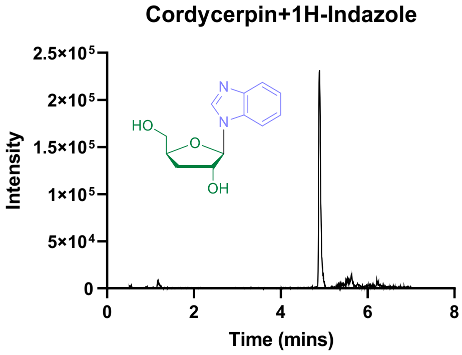

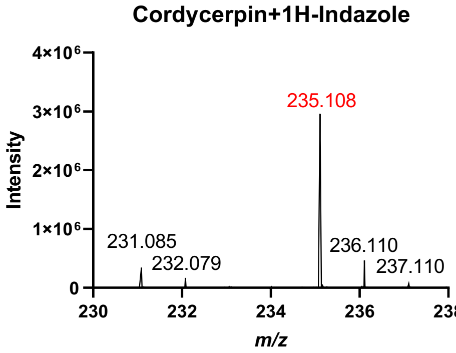

**Figure S1: Data for reactions to produce nucleosides.**

Product 3'-deoxy-15

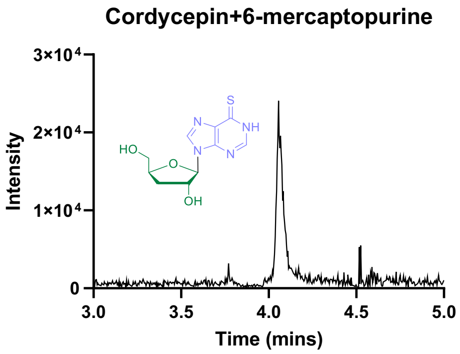

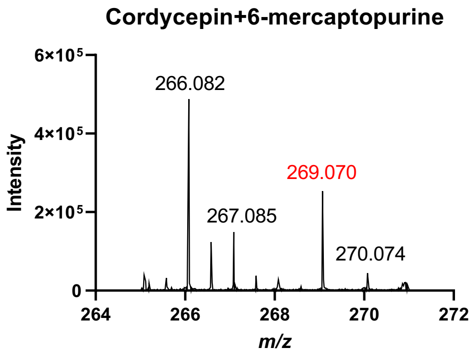

Product 3'-deoxy-38

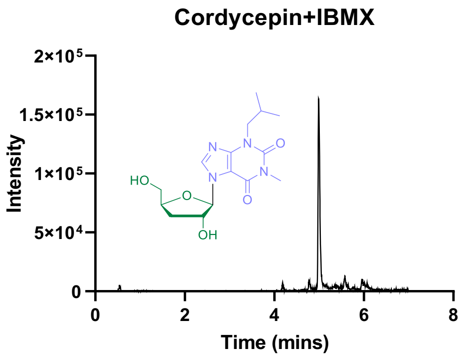

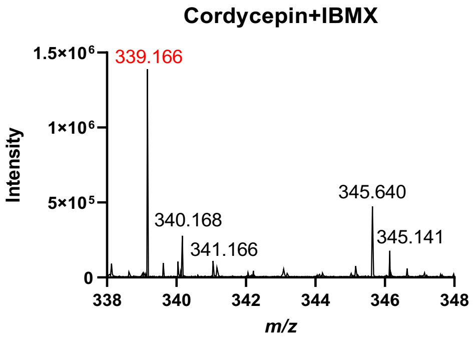

Product 3'-deoxy-54

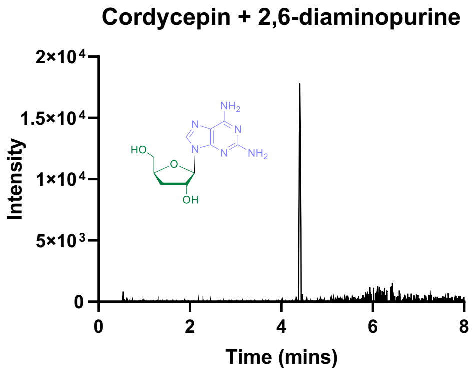

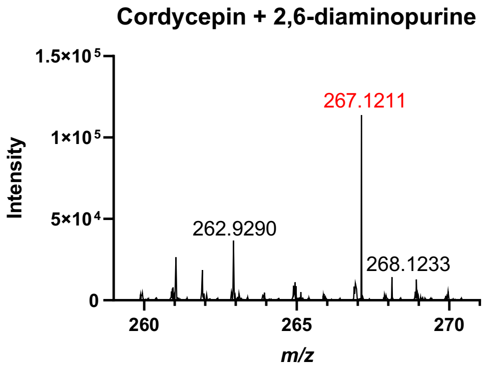

Product ribo-1

#### Figure S1: Data for reactions to produce nucleosides.

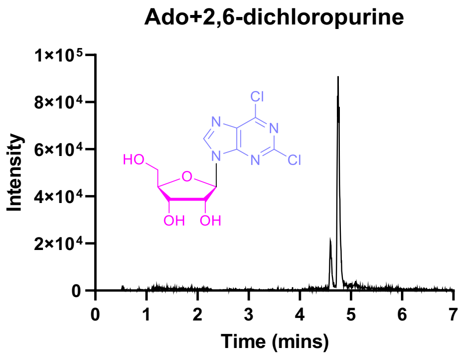

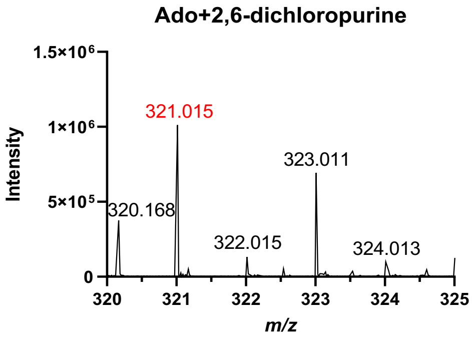

Product ribo-2

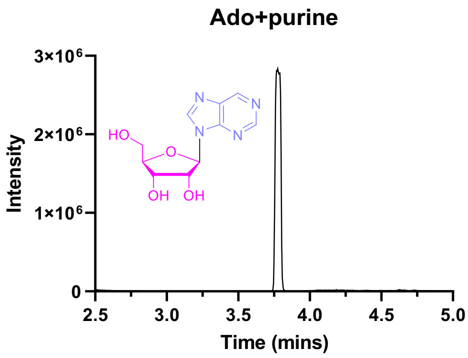

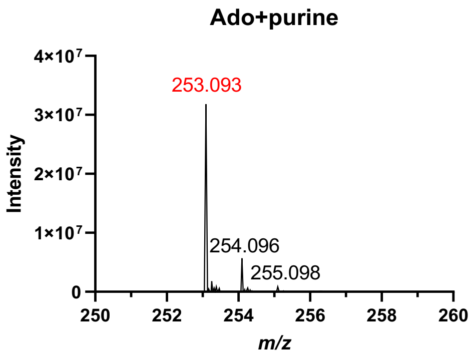

Product ribo-3

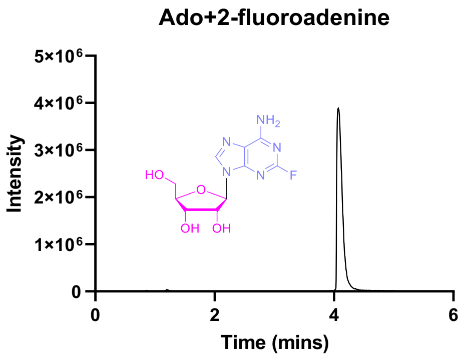

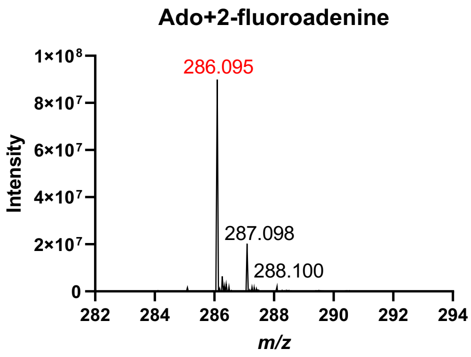

Product ribo-5

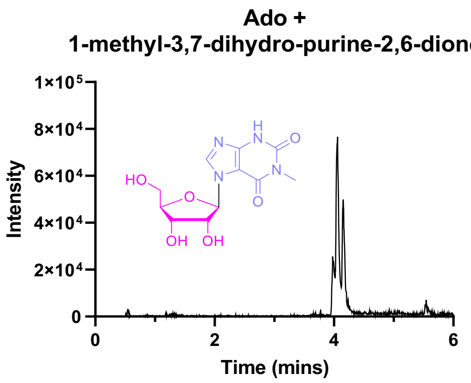

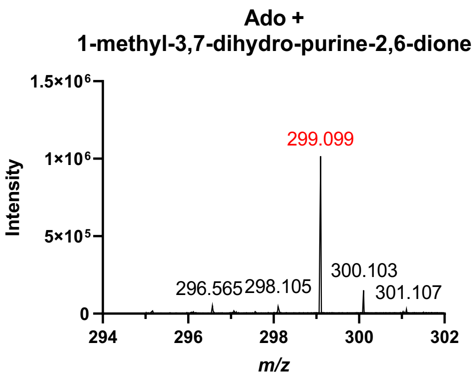

#### Figure S1: Data for reactions to produce nucleosides.

Product ribo-6

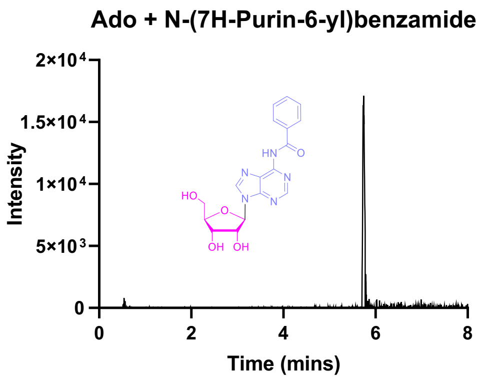

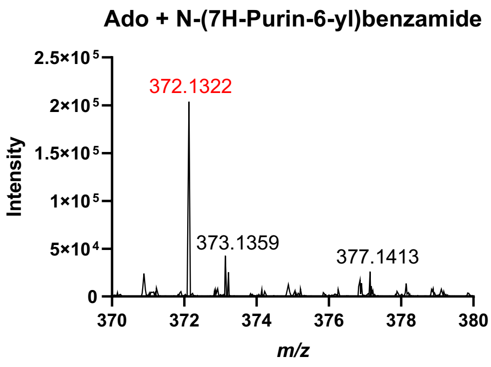

Product ribo-7

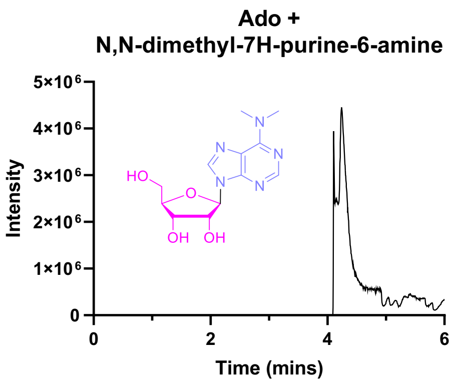

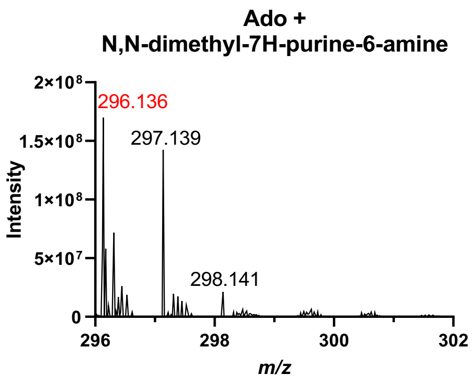

Product ribo-9

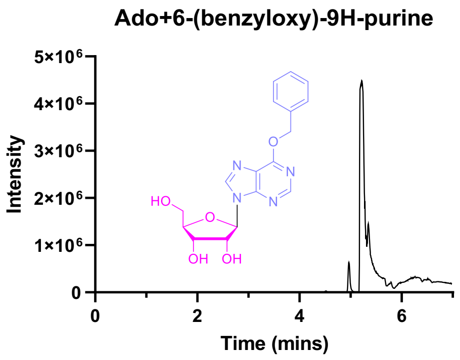

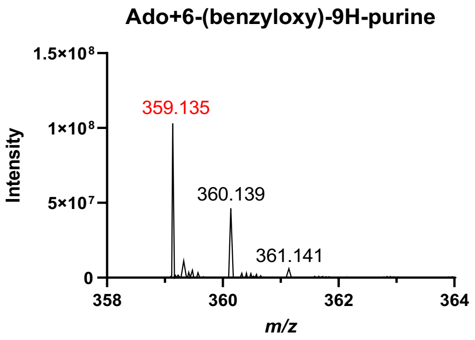

Product ribo-10

#### Figure S1: Data for reactions to produce nucleosides.

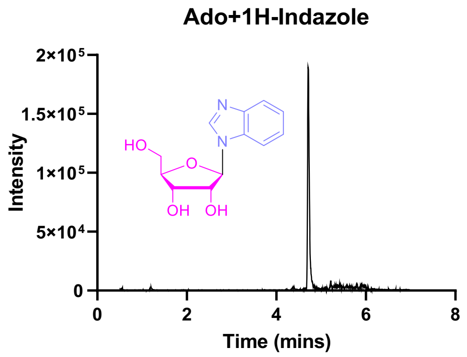

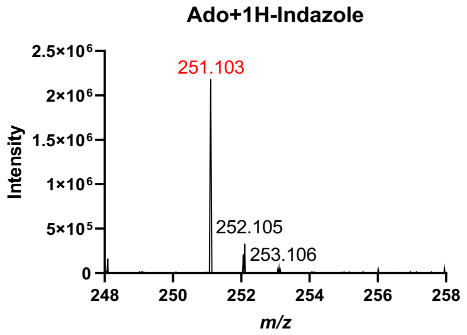

Product ribo-13

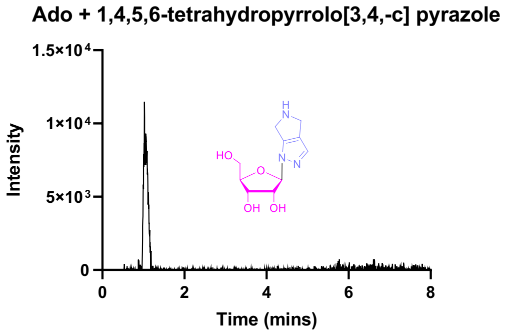

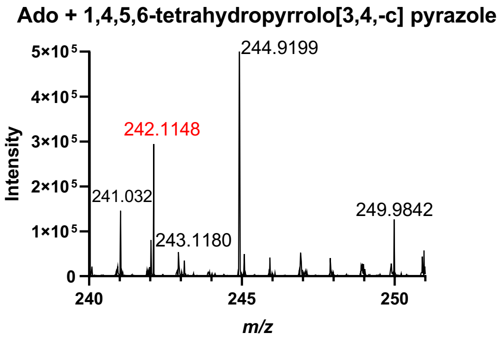

Product ribo-14

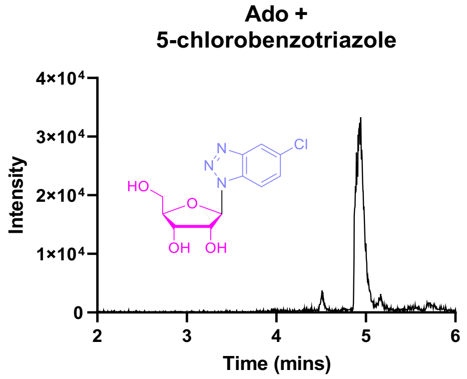

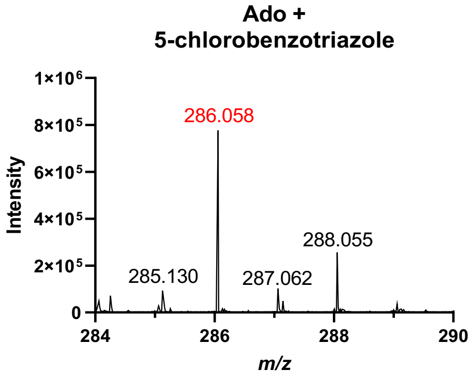

Product ribo-15

#### Figure S1: Data for reactions to produce nucleosides.

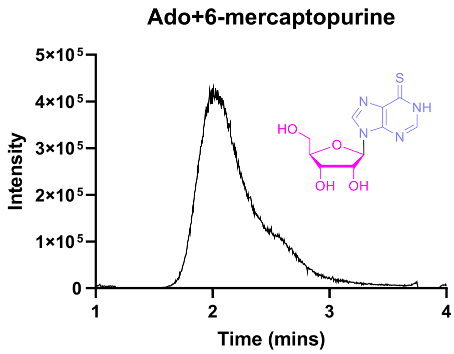

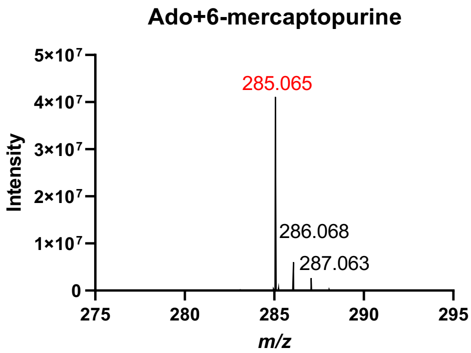

Product ribo-38

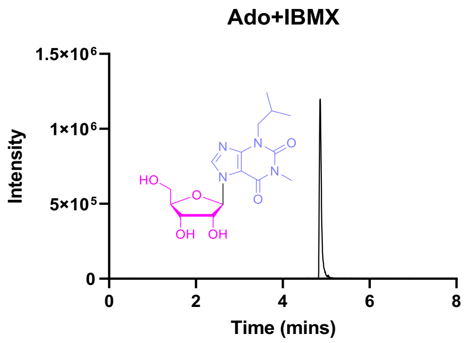

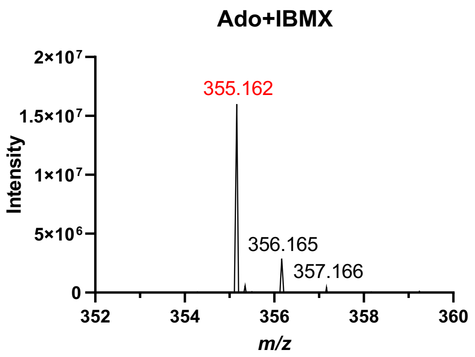

Product ribo-47

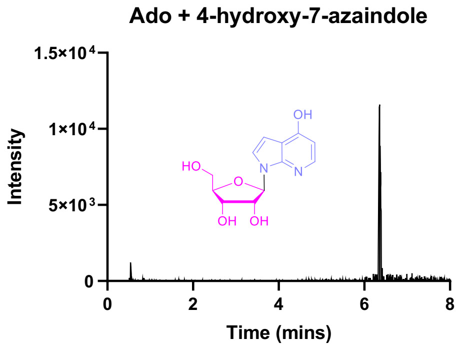

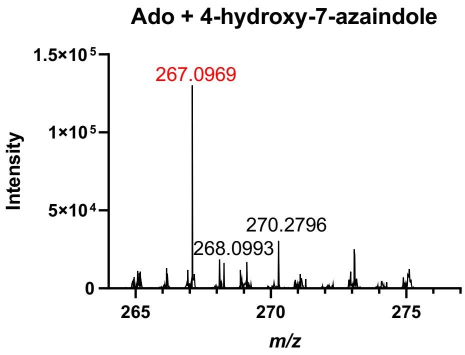

Product ribo-54

### Figure S1: Data for reactions to produce nucleosides.

**Ado + 2,6-diaminopurine**

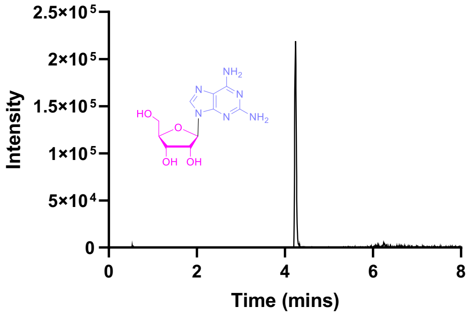

**Ado + 2,6-diaminopurine**

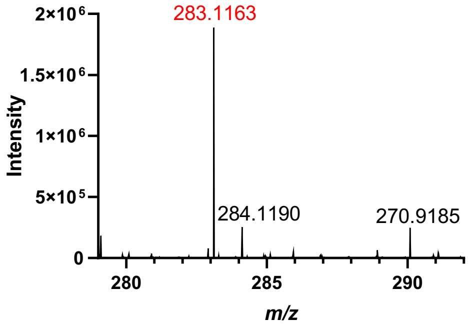

#### BpNDT

Product 2’-deoxy-18

**2'dGua+5-fluorouracil**

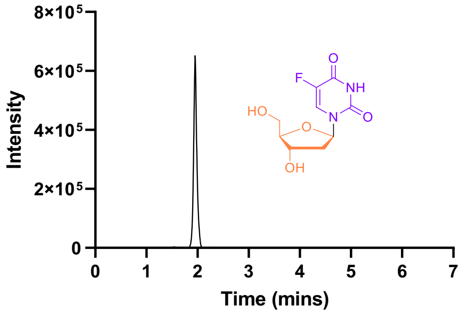

**2'dGua+5-fluorouracil**

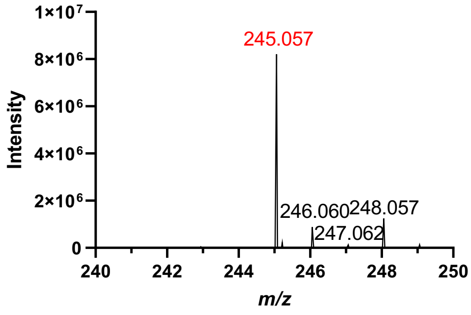

Product 2’-deoxy-20

**2'dGua+isocytosine**

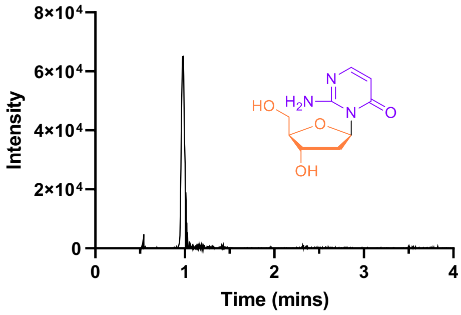

**2'dGua+isocytosine**

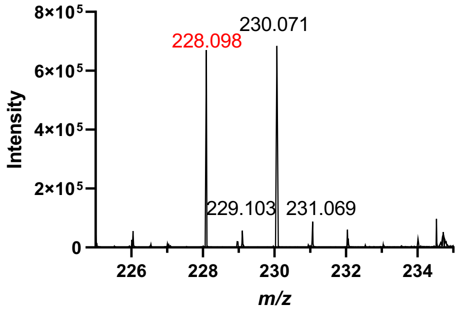

Product 2’-deoxy-21

**2'dGua+5-azacytosine**

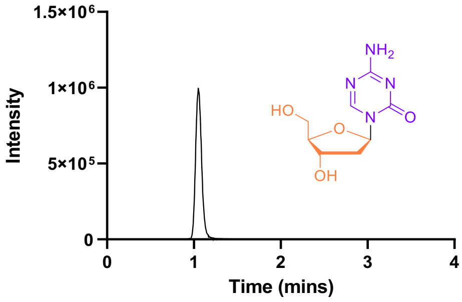

**2'dGua+5-azacytosine**

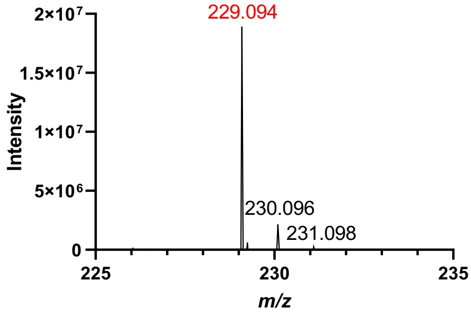

#### Figure S1: Data for reactions to produce nucleosides.

Product 2’-deoxy-23

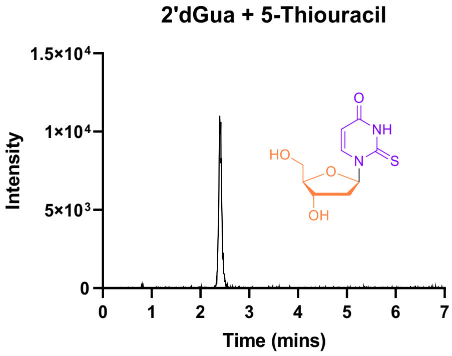

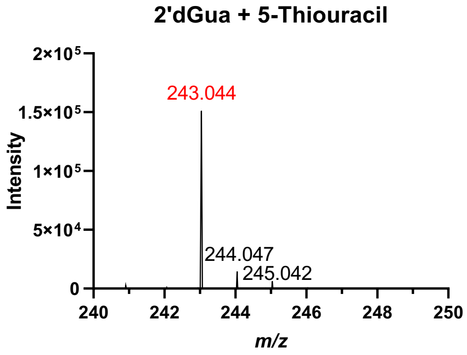

Product 2’-deoxy-24

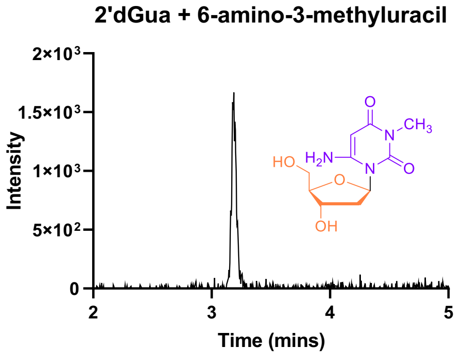

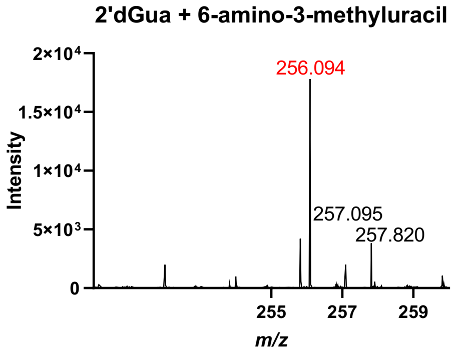

Product 2’-deoxy-26

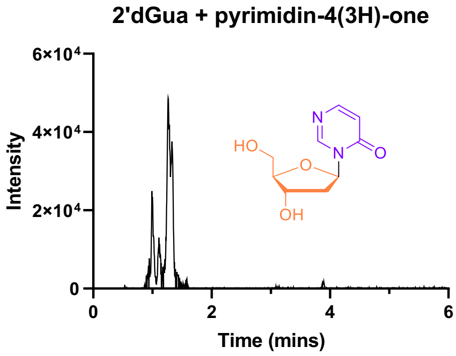

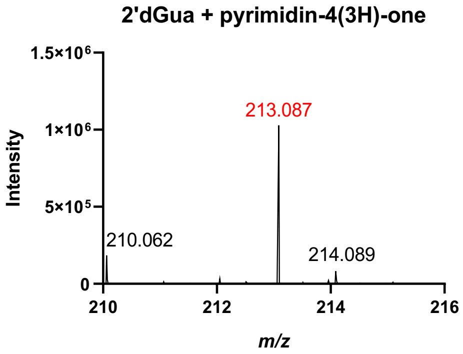

Product 2’-deoxy-29

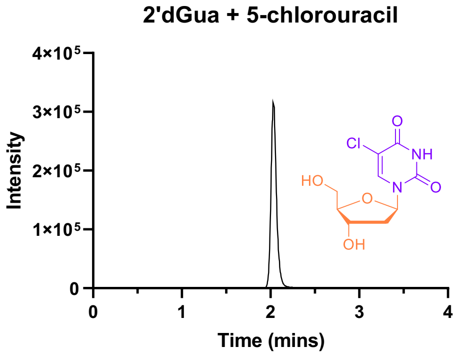

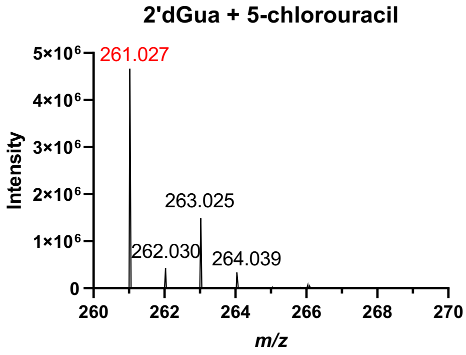

Product 2’-deoxy-30

#### Figure S1: Data for reactions to produce nucleosides.

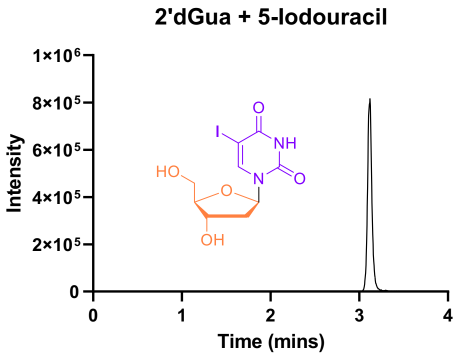

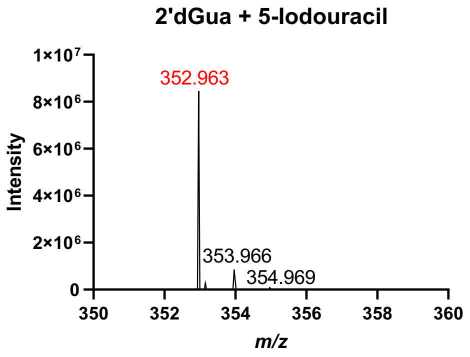

Product 2'-deoxy-32

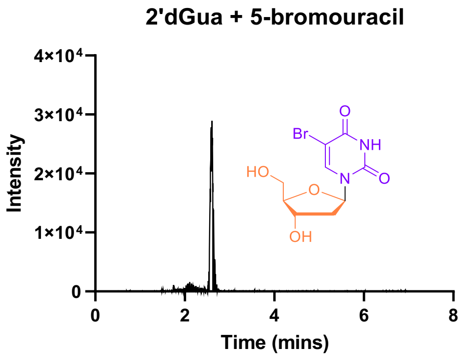

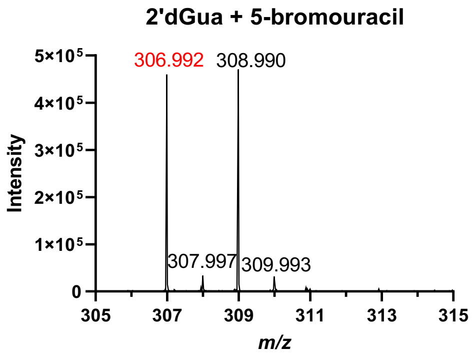

Product 2'-deoxy-43

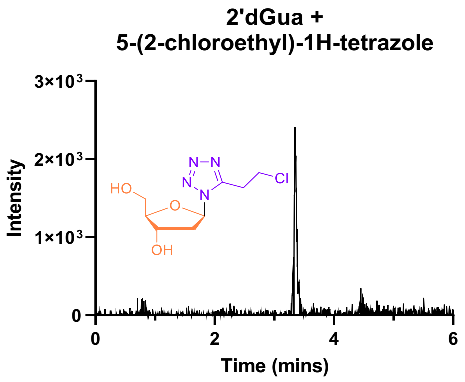

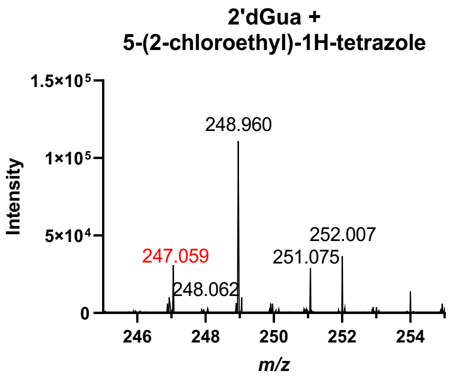

### BpNDT Y5F

Product ribo-18

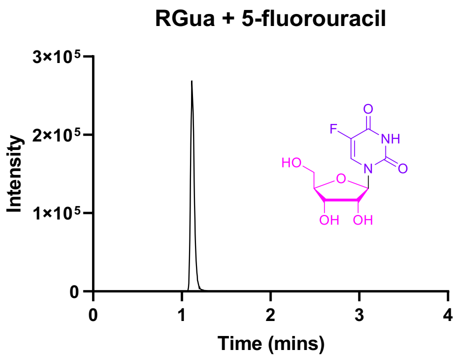

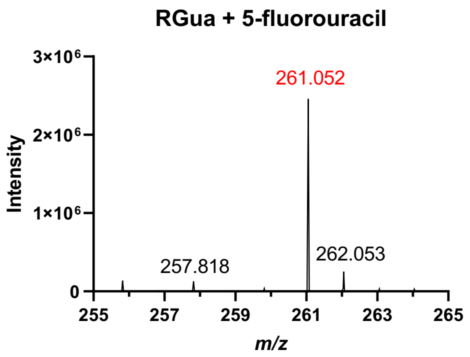

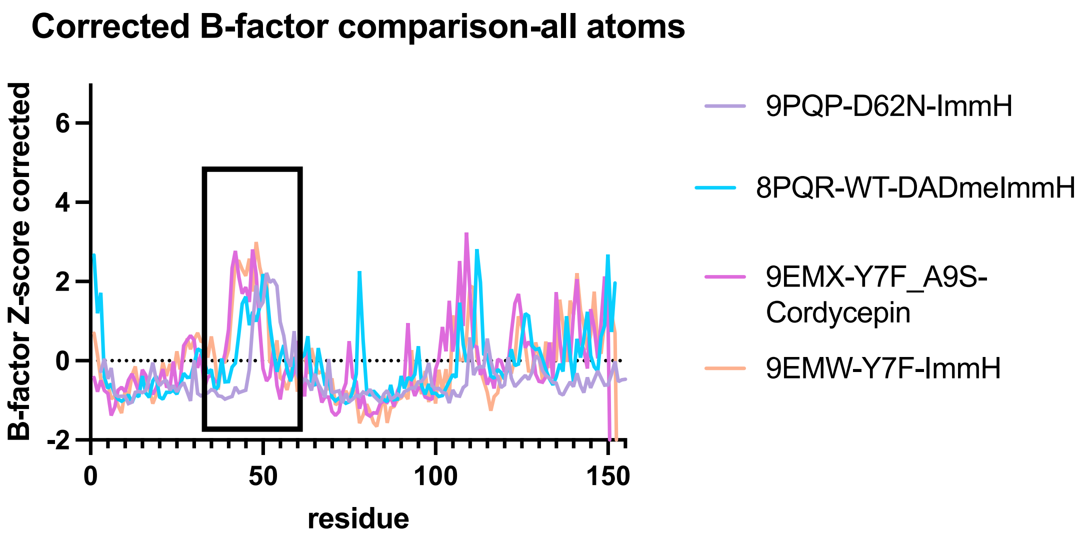

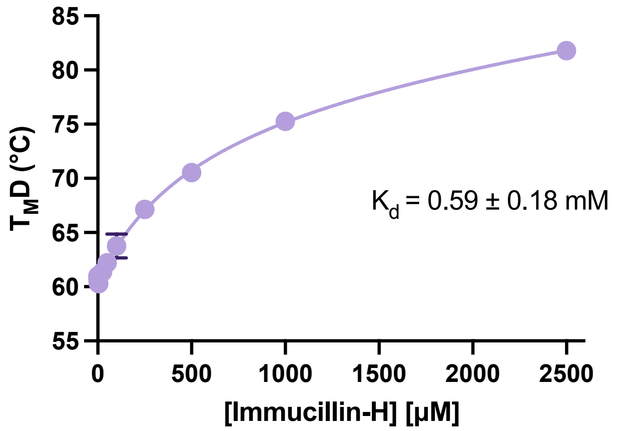

**Figure S2: Interactions of *Ct*NDT with transition state analogue DADmeImmH and CtNDT$_{Y7F\text{-}A9S}$ with substrate analogue ImmH.** Top: Comparison between apo and CtNDT bound to transition state analogue DADmeImmH1, depicting a region of higher B-factors in the apo enzyme which undergoes a small degree of stabilization upon tight binding of this analogue. Corrected B-factors ploted were obtained using Baverage from the CCP4 suite[6] and the correction previously described[7]. The correction is needed to compare B-factors as they can be affected by resolution and refinement strategies. Bottom: Binding curve for *Ct*NDT$_{Y7F\text{-}A9S}$ and Immucillin-H, carried out using differential scanning fluorimetry.

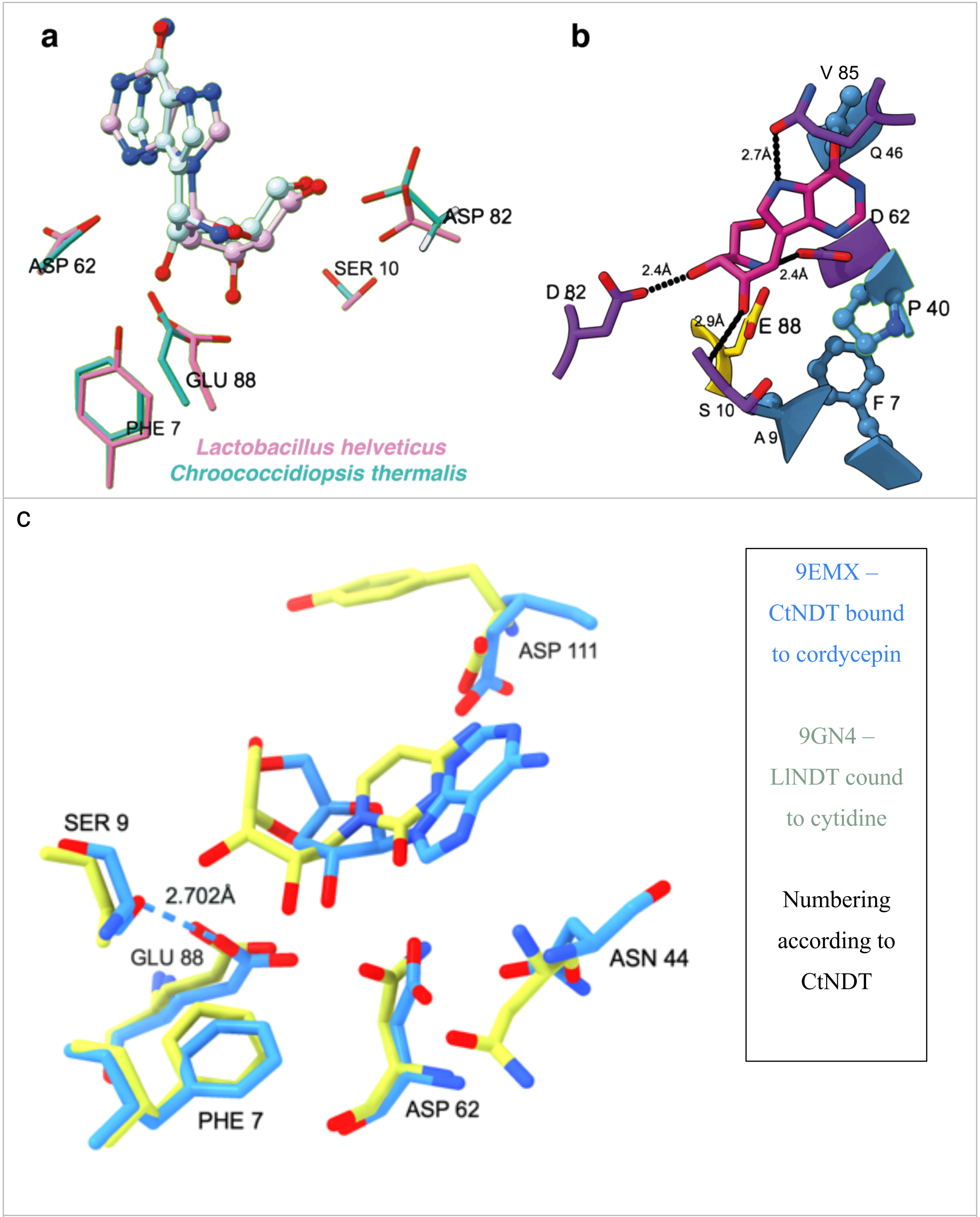

d

| | | Area (SA) Å2 | Volume (SA) Å3 |
|---|---|---|---|
| Enzyme | pdb | Pocket area | Pocket volume |
| *Chroococcidiopsis thermalis* | 8PQQ | 251.93 | 149.949 |
| *Trypanosoma brucei* | 2A0K | 264.734 | 147.782 |
| *Lactobacillus leichmannii* | 1F8X | 148.517 | 170.618 |

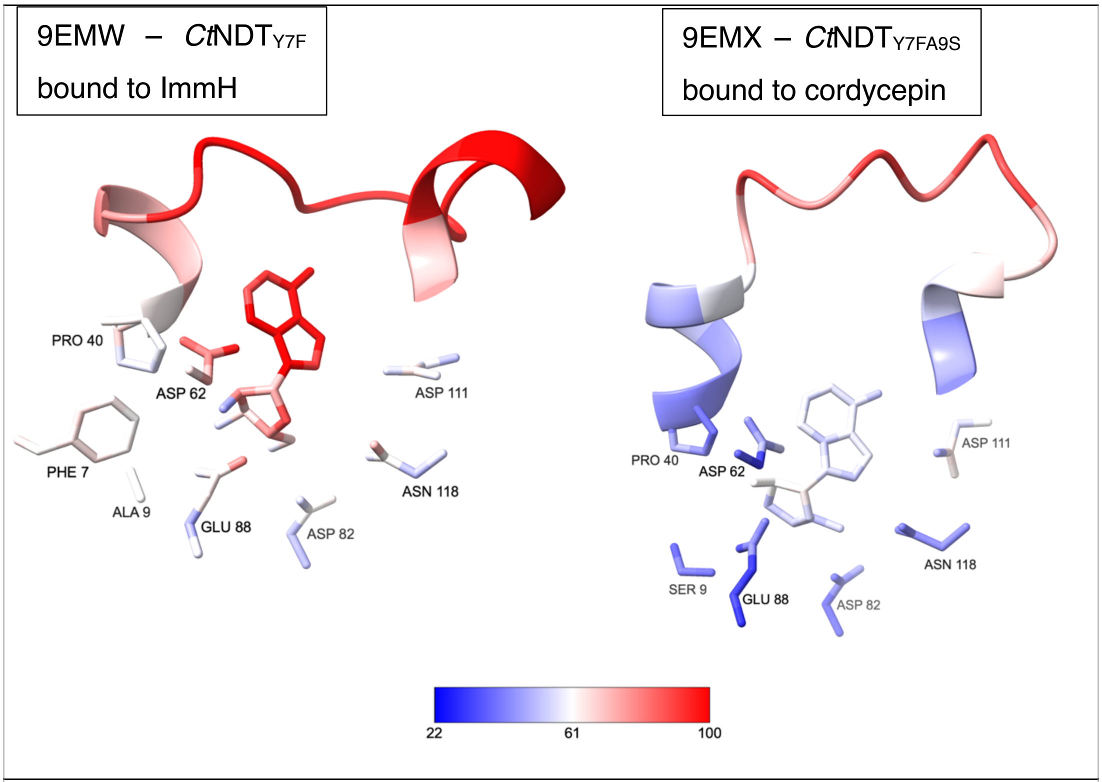

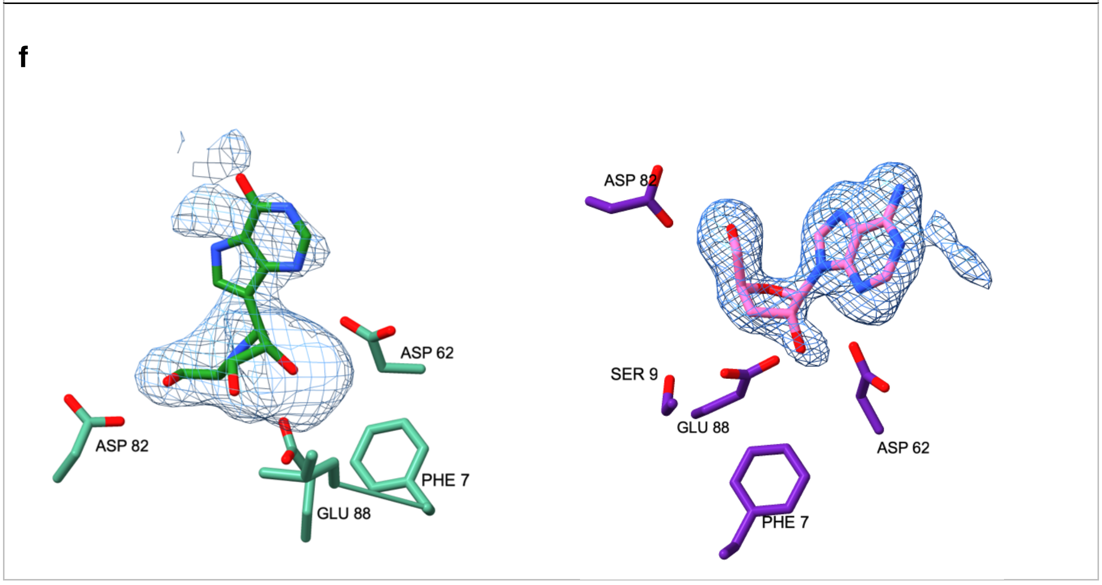

**Figure S3: Ligand binding site of NDT enzymes. a and b, residues interacting with ribosyl moiety of nucleoside susbtrates.** a) Comparison between *Lactobacillus helveticus* (pink, 1s2g[8], complex with 2'deoxyadenosine, rmsd 0.89 Å) and $CtNDT_{Y7F}$ (blue, this work, pdb 9EMW, complex with ImmH). b) Distances between key residues in the binding pocket for $CtNDT_{Y7F}$ interacting with ImmH. d) Comparison between substrate binding pocket volumes of different NDT enzymes. e) Residues and ligand atoms colored by normalized B factor (using Baverage -

https://www.ccp4.ac.uk/html/baverage.html), where smaller B factors are in blue and larger B factors are in red as per the color key in the bottom of the figure. f) Likelihood weighted mFo-DFc omit map at 2.5 σ contour (generated using Phenix maps), left 9EMW – *Ct*NDT$_{Y7F}$ bound to ImmH, right 9EMX – *Ct*NDT$_{Y7FA9S}$ bound to cordycepin).

### Figure S4. HPLC and raw kinetic data for all enzyme variants tested.

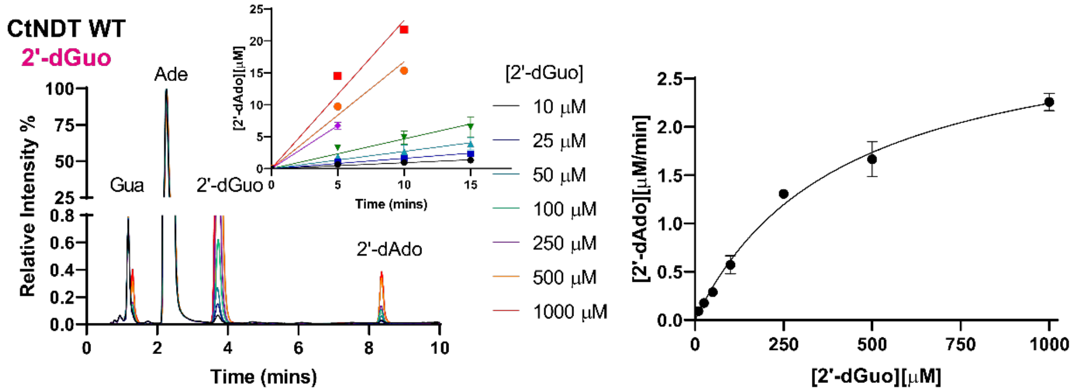

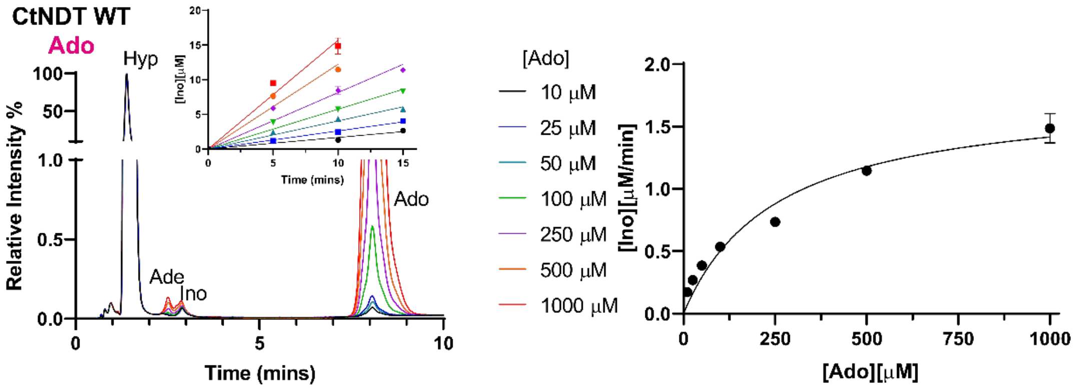

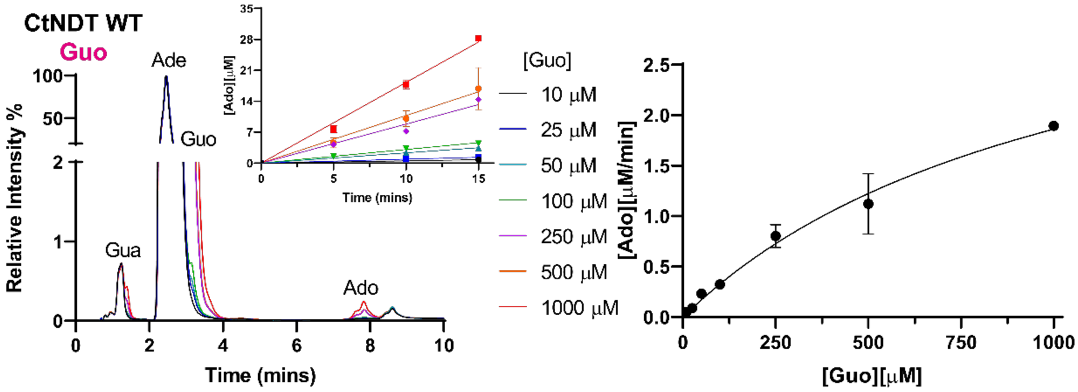

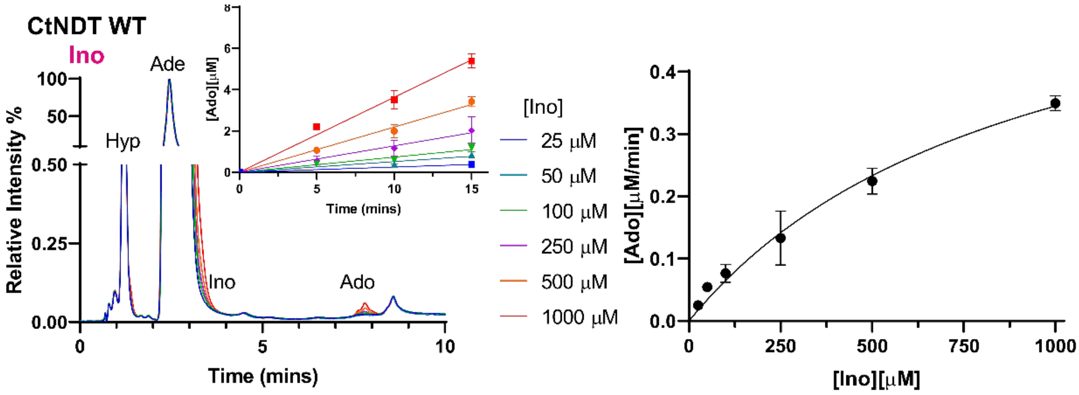

#### Figure S4. HPLC and raw kinetic data for all enzyme variants tested.

CtNDT WT
Uric acid

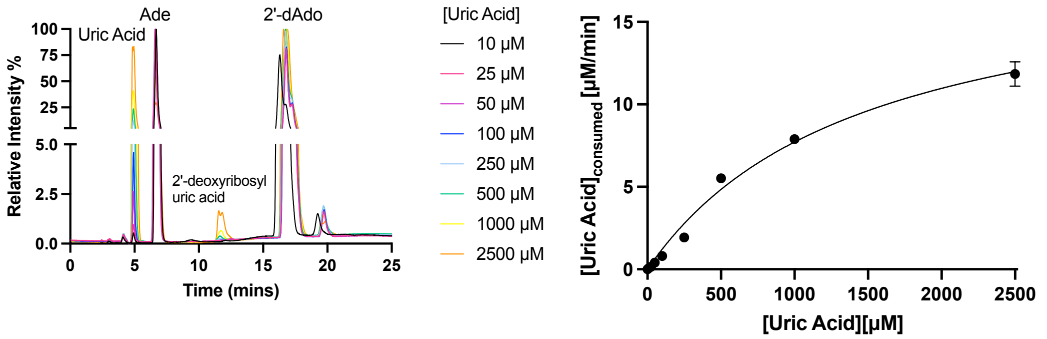

**CtNDT Y7F**
**2'-dGuo**

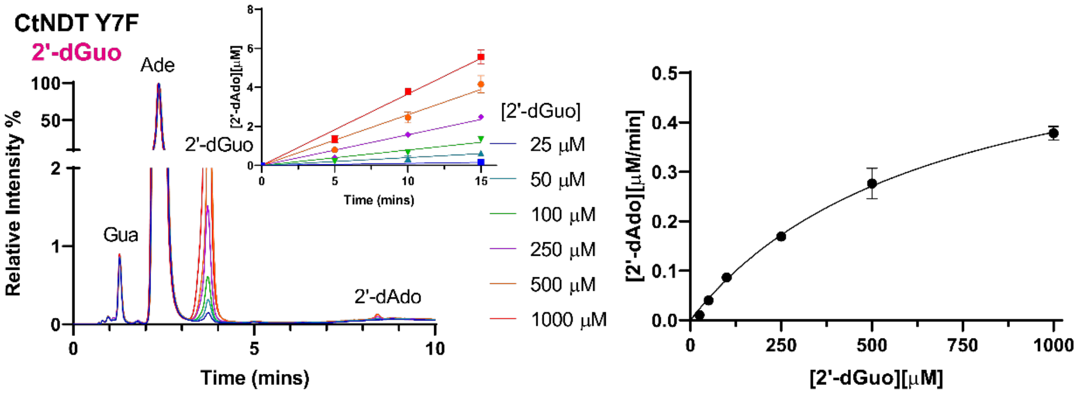

#### Figure S4. HPLC and raw kinetic data for all enzyme variants tested.

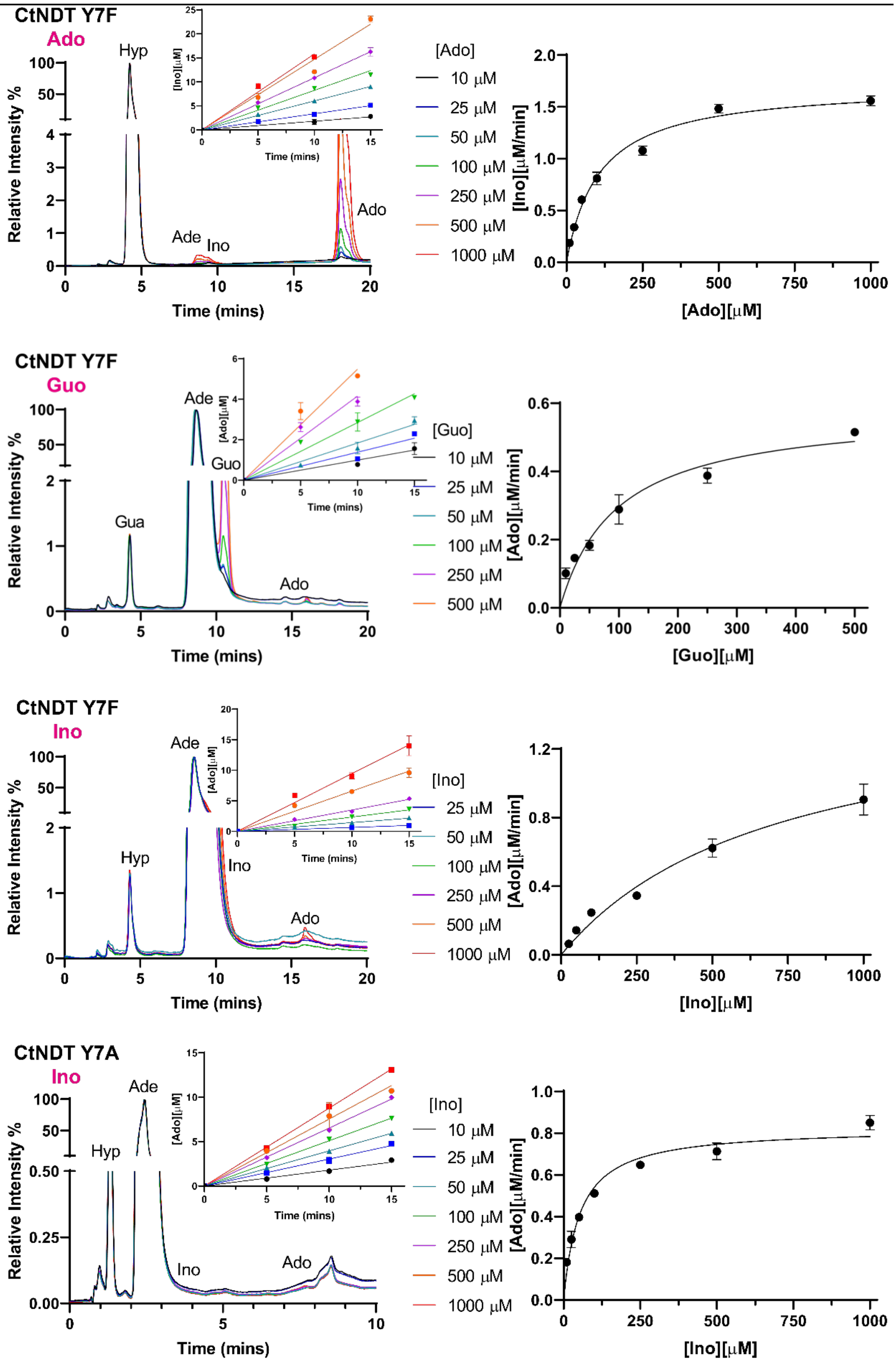

**Figure S4. HPLC and raw kinetic data for all enzyme variants tested.**

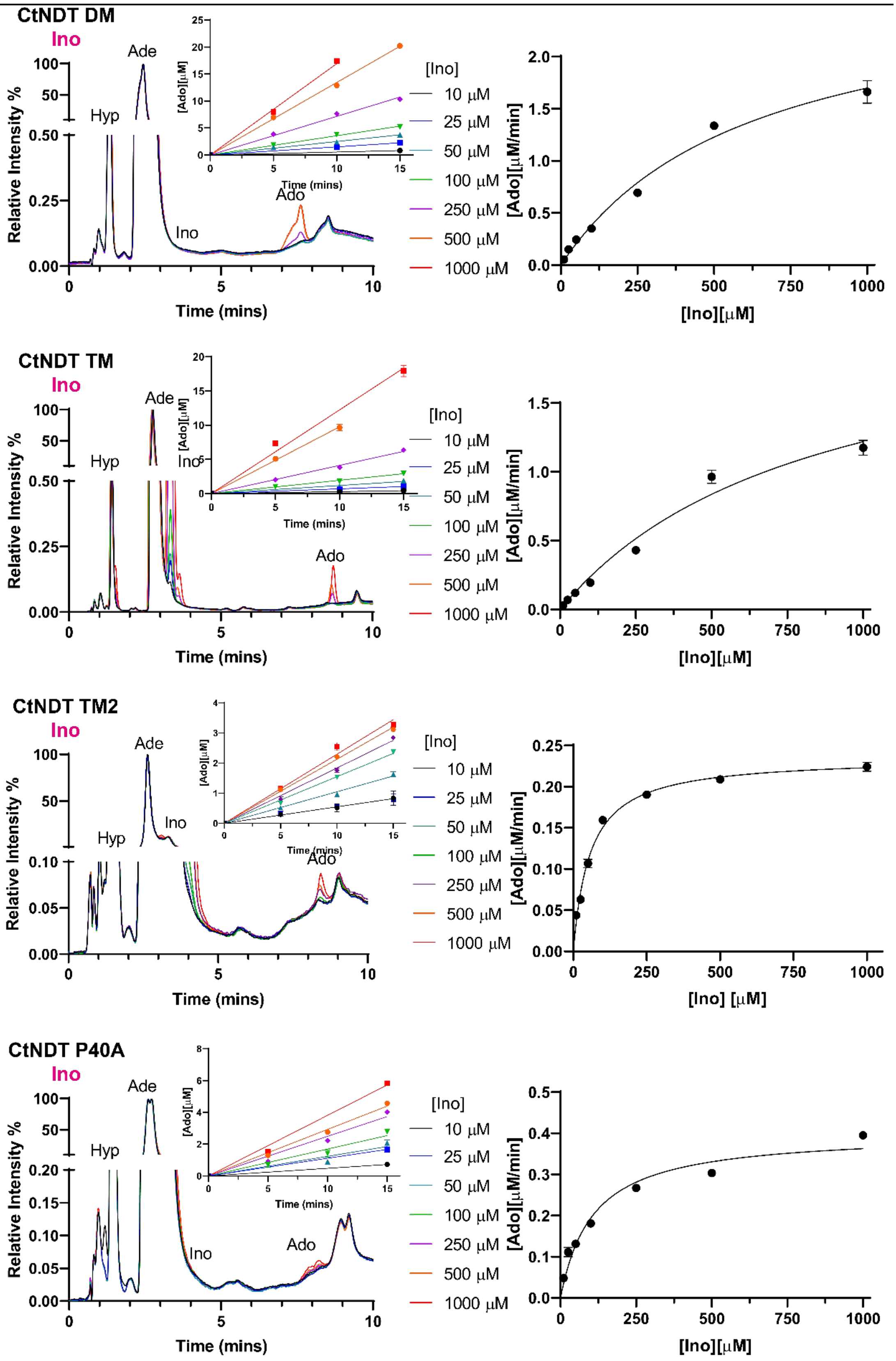

**Figure S4. HPLC and raw kinetic data for all enzyme variants tested.**

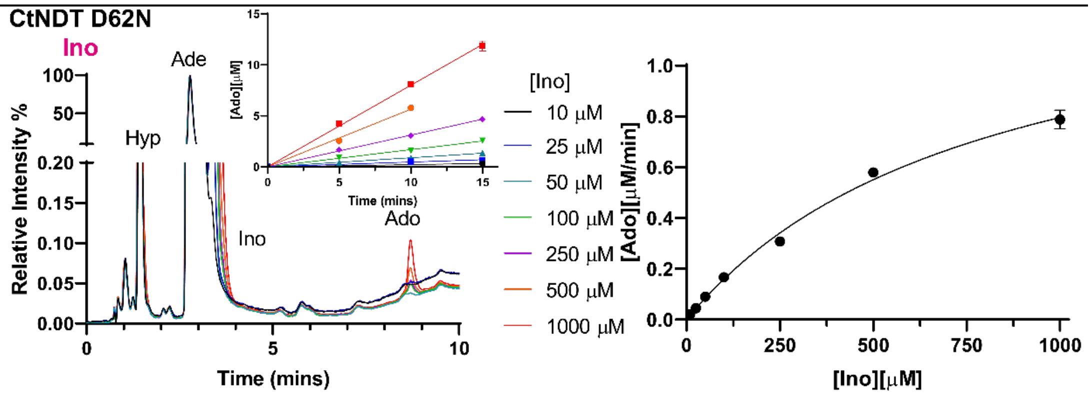

**Figure S4. HPLC and raw kinetic data for all enzyme variants tested.** Left, HPLC traces (260 nm absorbance), inset depicts replots of integrated area valus for each peak. A calibration curve with known quantities of Ado was used to convert are values into concentration. Right, replot of rates of product formation in function of substrate concentration to obtain kinetic parameters after fitting to a Michaelis-Menten equation.

**Figure S5: Proof of concept for nucleoside production using CtNDTY7F-A9S.** Our procedure generated nucleosides in a single step with the conditions shown below. Compound identities were confirmed by comparison to commercial standards of each new nucleoside product, and quantitified based on calibration curves with authentic standards.

a

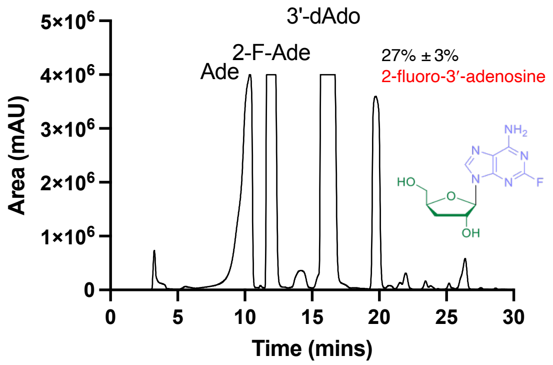

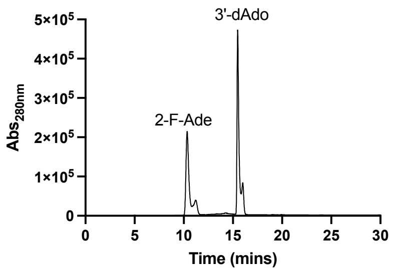

Left, HPLC trace (UV 260 nm, AU = arbitrary units) for purification of 2-fluoro-3′-deoxyadenosine (or 2F-3′d-Ado or 2F-cordycepin, Product 3’-deoxy-3). Reaction was carried out with 1:10 2-fluoroadenine to cordycepin substrate ratio. Quantifying 3′-deoxyadenosine decay reveals 27% of starting nucleoside substrate was hydrolysed. Right, reaction control with no enzyme added.

b

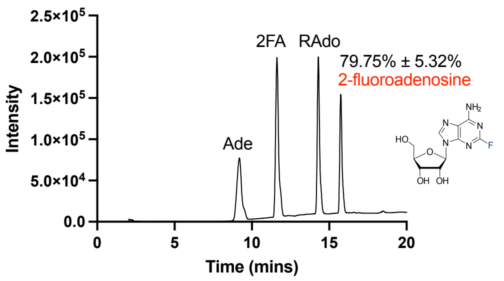

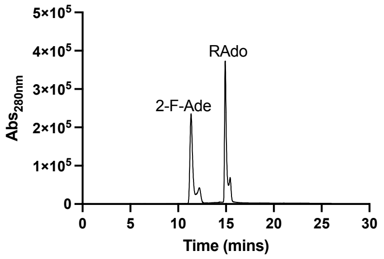

Left, HPLC trace (UV 260 nm)for purification of 2-fluoroadenosine (Product ribo-3). Quantifying adenosine decay reveals negligible hydrolysis. Right, reaction control with no enzyme added.

c

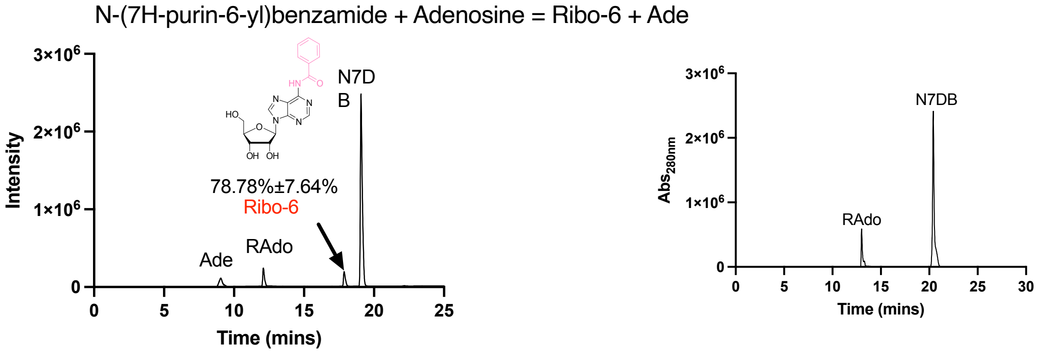

Left, HPLC trace (UV 260 nm) for purification of Product ribo-6. Reaction was carried out with 1:10 Adenosine to N-(9H-purin-6-yl)benzamide substrate ratio. Quantifying adenosine decay reveals negligible hydrolysis. Right, reaction control with no enzyme added.

d

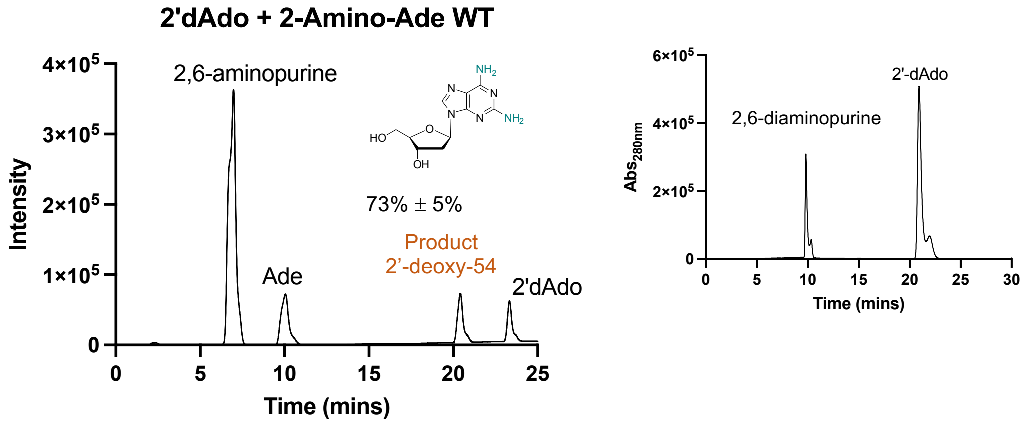

HPLC trace (UV 260 nm) for purification of Product 2'-deoxy-54 in a reaction with wild type enzyme. Reaction was carried out with 1:10 Adenosine to 2,6-diaminopurine substrate ratio. Quantifying 2′-deoxyadenosine decay reveals 15% hydrolysis. Right, reaction control with no enzyme added.

e

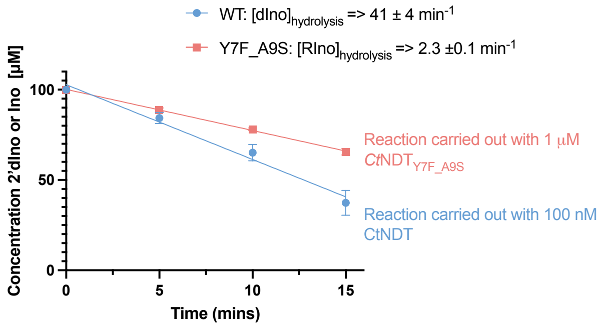

Time course for hydrolysis of 100 µM 2'-deoxyinosine using wild type *Ct*NDT or inosine using *Ct*NDT$_{Y7F_A9S}$ in the presence of 1mM 2,6-diaminopurine.

**Figure S6: Reaction tests to improve reaction yield and optimize nucleoside product formation**. a) varying pH while using 1:1 2'-deoxy-guanosine and adenine as substartes, reaction conducted for different amounts of time as indicated on graph. b) varying temperature while using 1:1 2'-deoxy-guanosine and adenine as substrates. c) testing whether adding xanthine oxidase to reactions in which the nucleoside donor is inosive changes equilibrium. Reaction with 1:1 or 1:10 2'deoxyinosine and adenine as substrates.

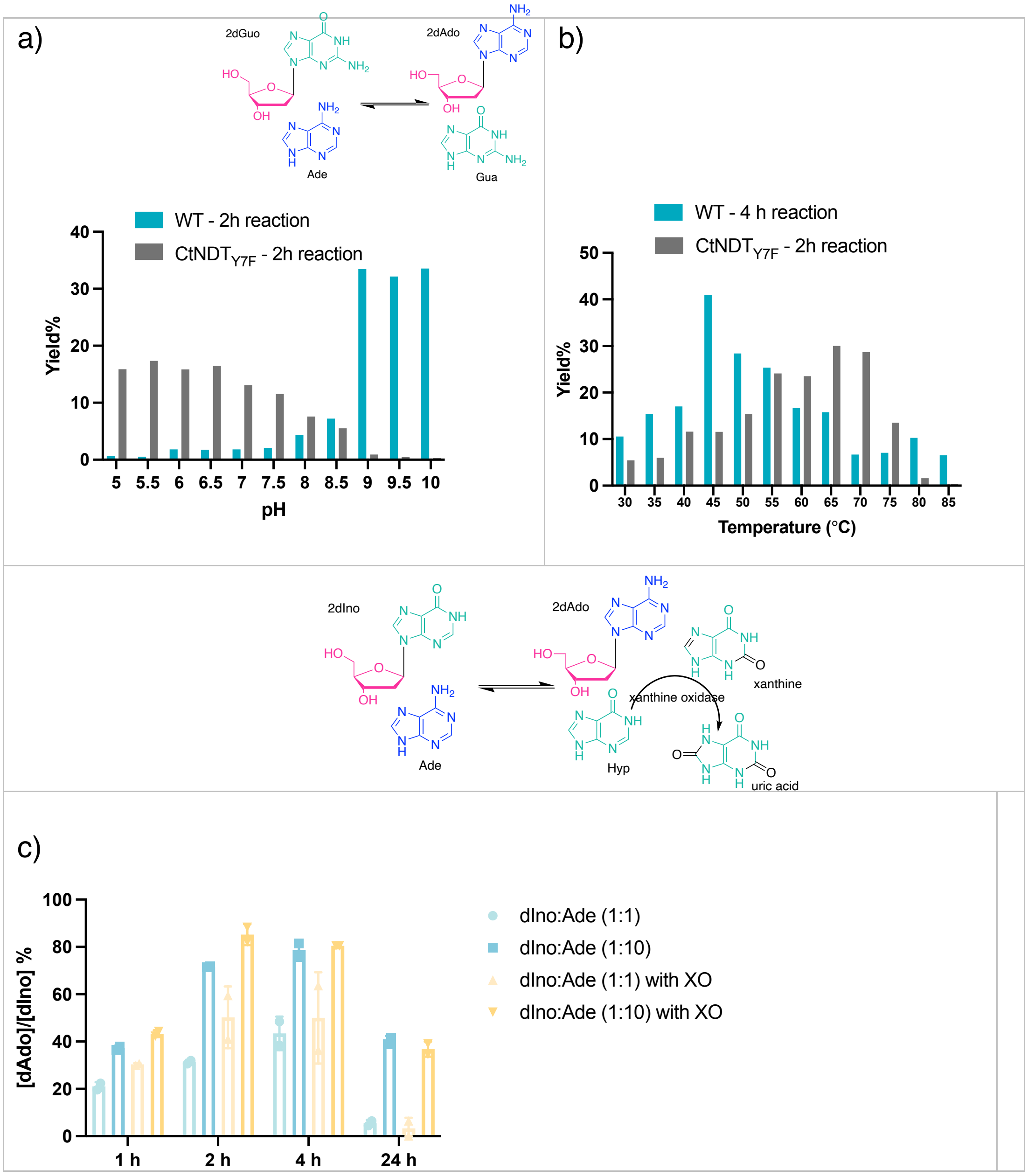

**Figure S7: SDS-PAGE for all proteins used in this work.** The orange boxes highlight fractions that were pooled and concentrated for use in subsequent experiments.

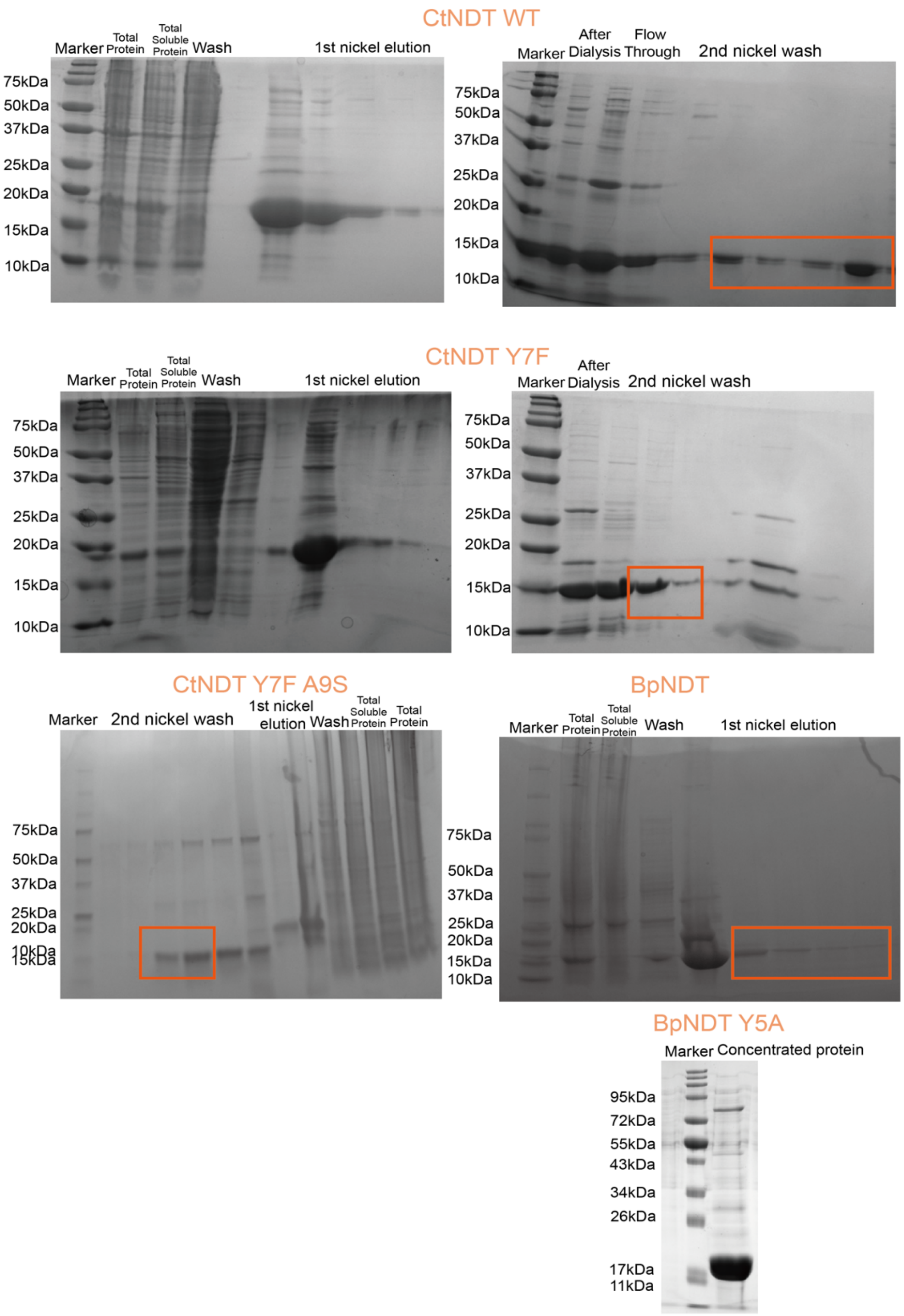

#### Figure S8: Intact protein mass spectrometry for all proteins employed in this work.

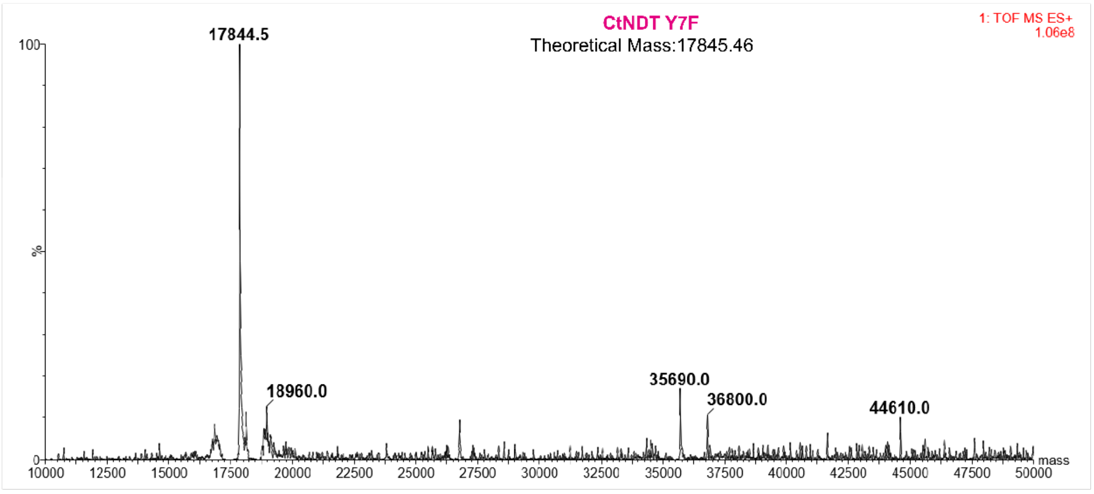

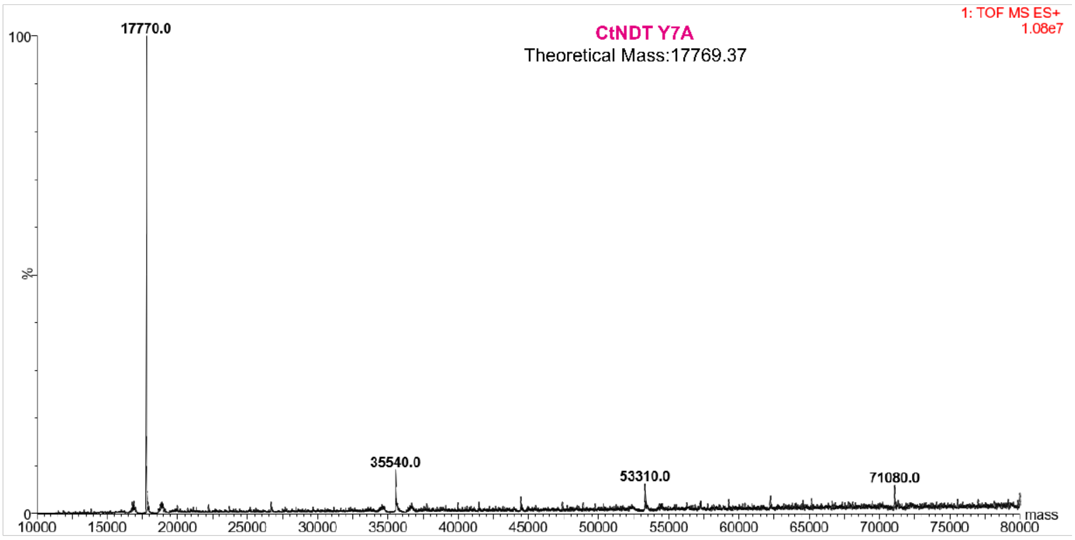

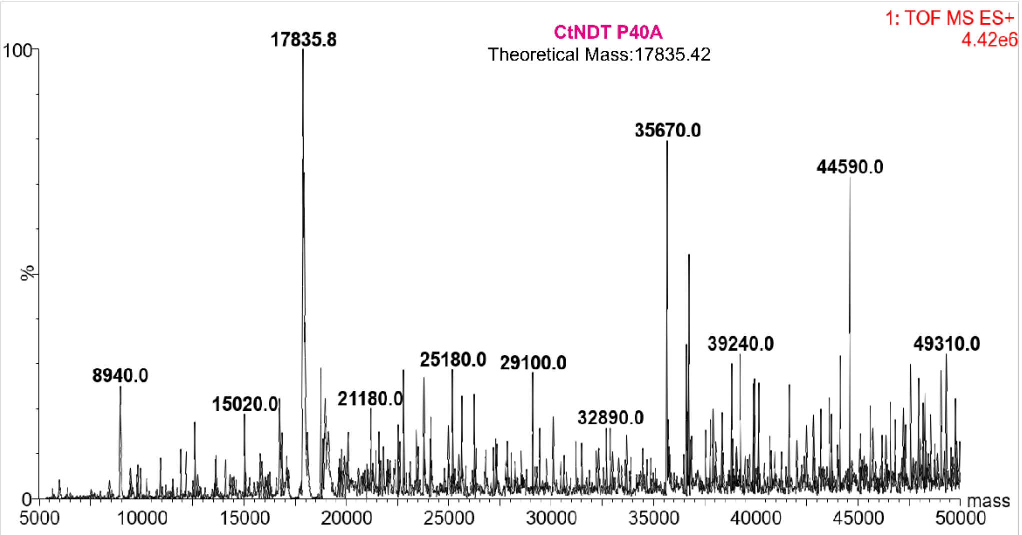

**Figure S8: Intact protein mass spectrometry for all proteins employed in this work.**

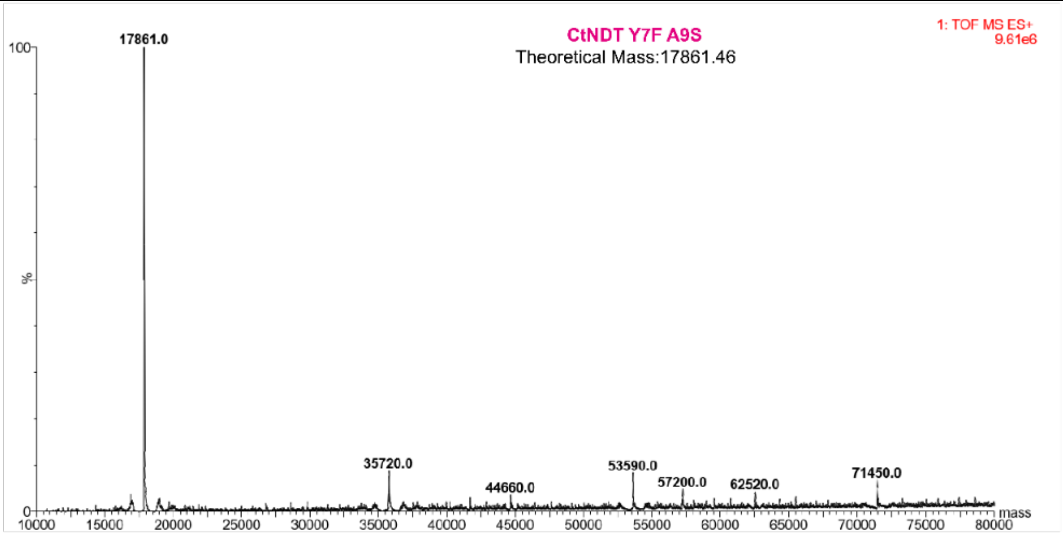

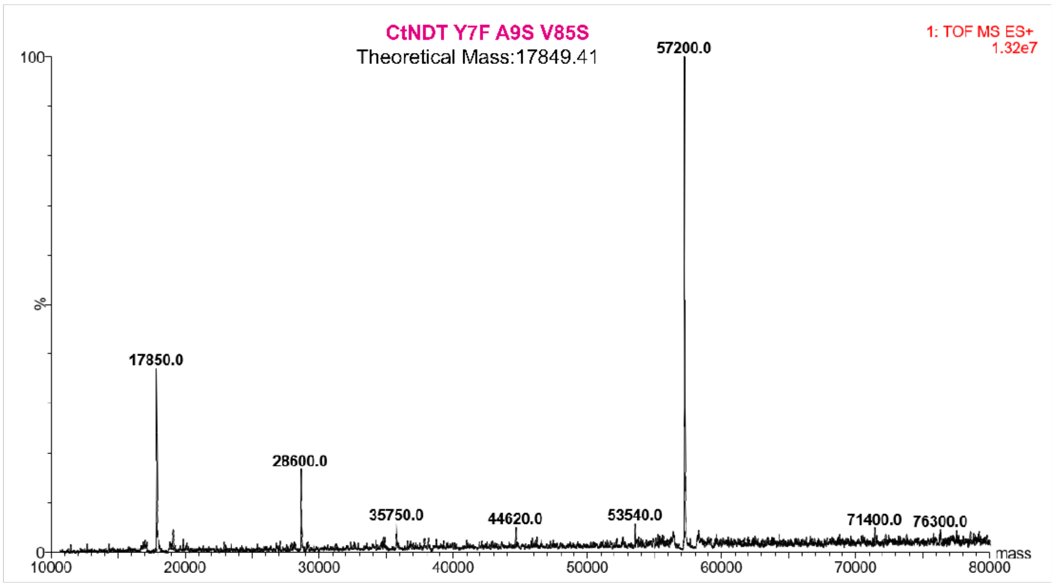

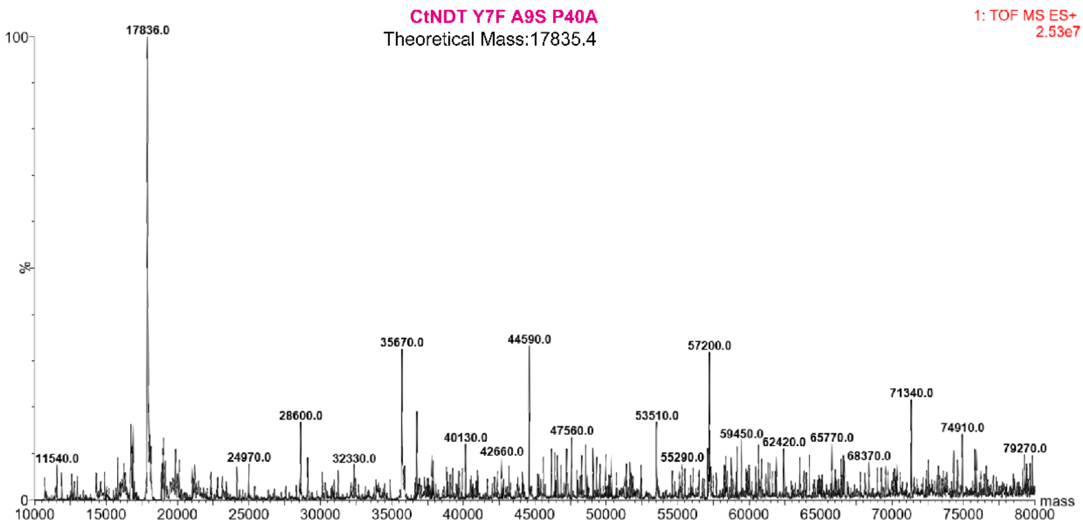

#### Figure S8: Intact protein mass spectrometry for all proteins employed in this work.

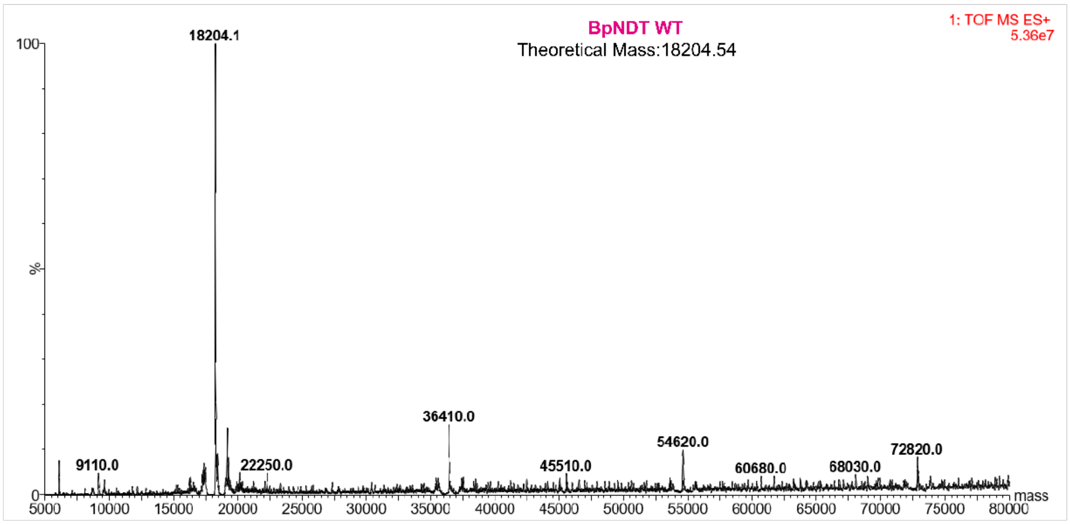

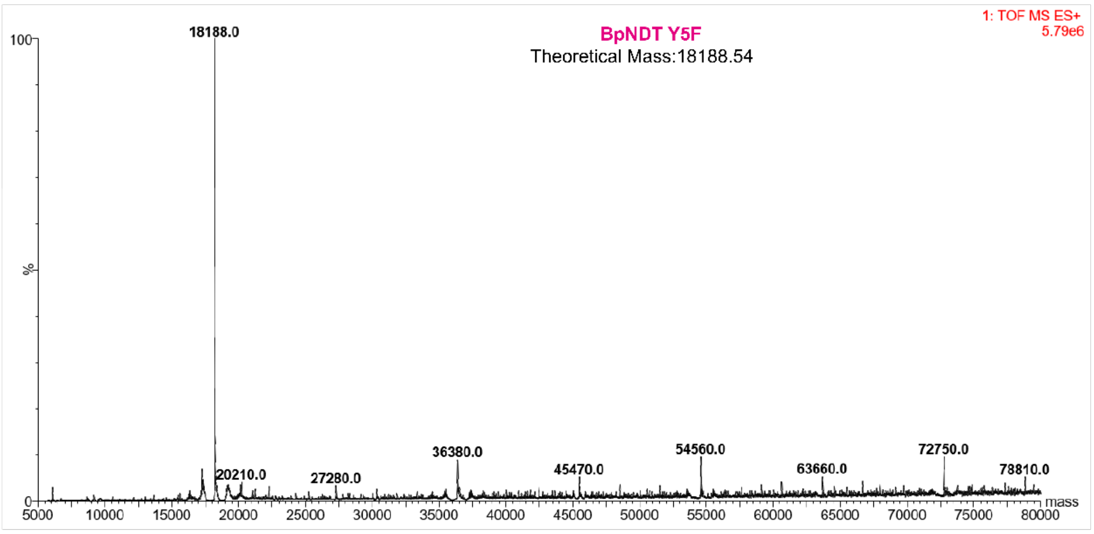

**Figure S8: Intact protein mass spectrometry for all proteins employed in this work.** Theoretical expected masses are shown as well as raw data.
